# BrumiR: A toolkit for *de novo* discovery of microRNAs from sRNA-seq data

**DOI:** 10.1101/2020.08.07.240689

**Authors:** Carol Moraga, Evelyn Sanchez, Mariana Galvão Ferrarini, Rodrigo A. Gutierrez, Elena A. Vidal, Marie-France Sagot

## Abstract

MicroRNAs (miRNAs) are small non-coding RNAs that are key players in the regulation of gene expression. In the last decade, with the increasing accessibility of high-throughput sequencing technologies, different methods have been developed to identify miRNAs, most of which rely on pre-existing reference genomes. However, when a reference genome is absent or is not of high quality, such identification becomes more difficult. In this context, we developed BrumiR, an algorithm that is able to discover miRNAs directly and exclusively from sRNA-seq data. We benchmarked BrumiR with datasets encompassing animal and plant species using real and simulated sRNA-seq experiments. The results demonstrate that BrumiR reaches the highest recall for miRNA discovery, while at the same time being much faster and more efficient than the state-of-the-art tools evaluated. The latter allows BrumiR to analyze a large number of sRNA-seq experiments, from plants or animals species. Moreover, BrumiR detects additional information regarding other expressed sequences (sRNAs, isomiRs, etc.), thus maximizing the biological insight gained from sRNA-seq experiments. Finally, when a reference genome is available, BrumiR provides a new mapping tool (BrumiR2ref) that performs an *a posteriori* exhaustive search to identify the precursor sequences. The code of BrumiR is freely available at https://github.com/camoragaq/BrumiR.

## Introduction

MicroRNAs (henceforth denoted by miRNAs) are small RNA molecules usually shorter than 25 nucleotides (nt), which have been identified as crucial regulators of gene expression mostly at the post-transcriptional level (1). miRNAs are involved in a wide range of biological processes including cell cycle, differentiation, apoptosis and disease (2). They have been the target molecules for a large number of important applications, more particularly in cancer where miRNAs have been shown to play important roles in driving or suppressing tumor spread (3,4). In plant species, unravelling host-pathogen interactions mediated by miRNAs may shed light on plant development and its relation with the environment, both essential knowledge that can lead to the discovery of new biotechnological products for the agricultural industry (5,6).

Since the first classification and annotation of miRNAs (7,8), accurately identifying them as well as the regulatory networks in which they are involved has proven difficult (9,10). Accurate prediction of known and novel miRNAs along with their targets is however essential for increasing our understanding of the miRNA biology (3).

Nowadays, a common experimental practice is to identify miRNAs and their expression patterns using next generation sequencing technologies (NGS) (11). Commonly, NGS experiments are able to generate more than 20 million sRNA-seq reads, thus promoting the development of algorithms to transform and process such data into biological information (12).

Currently, there are two computational strategies for the discovery of miRNAs: 1) genome-based approaches that rely on the mapping of the sRNA-seq reads to a reference genome and subsequent evaluation of the sequences generating the characteristic hairpin structure of miRNA precursors (9); 2) machine-learning approaches which rely on the biogenesis features extracted from the knowledge on miRNA sequences available in databases such as miRBase (13) and on the analysis of the duplex structure of miRNAs (14). Genome-based methods, that have been updated at the pace of the evolving NGS technologies, are the most widely used tools in this field, and their results have populated the public miRNA repositories (12). Such methods are the natural choice for the study of model species with high quality reference genomes available. However, it has been shown that most of the genome-based tools struggle with a high rate of false positive predictions (9). Additionally, a critical step of such tools is the use of genome aligners (15,16) to map the sRNA-seq reads to the reference genome. Mapping short (< 30 nt) and very similar sequences to a large, complex, and repetitive reference genome is however a difficult and error-prone task (17). Genome-based methods are thus highly sensitive to the aligner selected as well as to the parameters employed and the thresholds chosen (*e.g.* number of mismatches allowed) in order to discard mapping artefacts generated from sequencing errors (18). Furthermore, despite all the advancements in the sequencing technologies and *de novo* assembly methods, few complete genomes are available today, which is a recurring problem that researchers working on non-model species face (19). The lack of a high quality reference genome thus reduces the possibilities for discovering novel miRNAs (14). Genome-based methods such as miRDeep (20), miRDeep2 (21), and miR-PREFeR (22) are included in this group.

On the other hand, new methods such as miReader (23), MirPlex (24), and mirnovo (14), in particular using machine-learning approaches, were specifically developed as an alternative to discover miRNAs in species without a reference genome. In the case of mirnovo, the initial step involves the clustering of the sRNA-seq reads performing an all-vs-all read comparison that is followed by a subsequent classification of the clusters into putative miRNAs using pre-trained models. The performance obtained by such methods on well-annotated species is comparable to those achieved by genome-based methods (9). However, relying exclusively on annotated miRNAs for training machine learning models may introduce a bias towards the identification of well-characterized miRNAs over species-specific ones (12). Nonetheless, machine learning methods have demonstrated that it is possible to discover miRNAs using only the sequence information present in the sRNA-seq experiment (14).

There remains however a need to go further in the development of algorithms for finding novel miRNAs in non-model species using only the sequence information. With this purpose in mind, the adoption of a special type of graphs called *de Bruijn* graphs may be considered. This is a widely used approach for the *de novo* reconstruction of genome or transcriptome sequences (25). It therefore appears to be a plausible option for organizing, clustering and assembling the sequence information present in sRNA-seq experiments. However, accommodating the de Bruijn graph approach for the discovery of miRNAs involves the development of new methods to address the specific characteristics of sRNA-seq data. Indeed, mature miRNA sequences are short (18-24 nt), thus limiting the overlap length for building a de Bruijn graph which in turn impacts the global topology by inducing tangled graph structures. Moreover, miRNAs captured in a sRNA-seq experiment have variable expression, from low (few reads) to highly expressed (thousands of reads), which may induce spurious graph connections that should be removed in order to isolate and detect both types of miRNAs. Finally, the sequencing errors present in sRNA-seq data further induce spurious connections and are harder to detect as compared to genomic data due to the variable expression and the shorter lengths of the miRNAs. Overall, using a de Bruijn graph to analyze sRNA-seq data and extract information from such data seems thus counterintuitive as mature miRNAs are captured full-length by the current NGS technologies. However, a de Bruijn graph has several interesting properties for the discovery of miRNAs, mainly due to the fact that it encodes all the sRNA-seq sequence information at once in a compact and connected representation (graph), without the need to perform an all-vs-all read comparison or mapping to a reference.

In this paper, we present BrumiR, a *de novo* algorithm based on a de Bruijn graph approach that is able to identify miRNAs directly and exclusively from sRNA-seq data. Unlike other state-of-the-art algorithms, BrumiR does not rely on a reference genome, on the availability of close phylogenetic species, or on conserved sequence information. Instead, BrumiR starts from a de Bruijn graph encoding all the reads and is able to directly identify putative miRNAs on the generated graph. BrumiR also removes sequencing errors and navigates inside the graph detecting putative miRNAs by considering several miRNA biogenesis properties (such as expression, length, topology in the graph). Along with miRNA discovery, BrumiR can also assemble and identify other types of small and long non-coding RNAs expressed within the sequencing data. Finally, when a reference genome is available, BrumiR provides a new mapping tool (BrumiR2ref) that performs an exhaustive search to identify and validate the precursor sequences.

We extensively benchmarked BrumiR on animal and plant species using simulated and real datasets. The benchmark results demonstrate that BrumiR is very sensitive, besides being the fastest tool, and its predictions were supported by the characteristic hairpin structure of miRNAs. Finally, we also applied BrumiR to the discovery of miRNAs of *Arabidopsis thaliana* and identified three novel high-confidence miRNAs involved in root development. These putative miRNAs were not discovered before by any other software, thereby showing the potential of using different approaches even in the case where high quality genomes are available. The code of BrumiR is freely available at https://github.com/camoragaq/BrumiR.

## RESULTS

### BrumiR discovers mature miRNAs directly from the sRNA-seq reads

The main idea behind BrumiR is that mature miRNAs can be discovered directly from the information contained in the sequenced sRNA-seq reads. To achieve this, BrumiR starts by building a de Bruijn graph (using *k*-mers of size 18) from the sRNA-seq reads, then compacting all the simple nodes thus leading to the unipath graph (27) (Figure 1.1, Methods section). The unipath graph encodes all the sequence information of the sRNA-seq experiment, including sequencing errors, adapters, and other types of sequences (Figure 1.1). The construction of the unipath graph allows to avoid entirely the alignment of the sRNA-seq reads to a reference genome. Following the unipath graph construction, BrumiR cleans the graph by removing tips (dead-end nodes) with low expression/abundance (KM < 5), which are usually generated from sequencing errors (Figure 1.2). One feature of the miRNA biogenesis is that after Dicer cleavage, the mature miRNA is the most abundant of the three by-products and when it is sequenced, it has a uniform expression along its sequence (20). Therefore, BrumiR expects that the neighbor elements within a particular putative miRNA will have similar expression. BrumiR checks all neighbor connections (arcs), and deletes any connection with a relative expression difference larger than 3 fold (Figure 1.3, Methods section), and the new graph is cleaned again by removing tips (Figure 1.4). Clusters of unipaths (connected components) with topologies related to sequencing errors are also removed (Figure 1.5, Methods section). BrumiR attempts to re-assemble all unipaths within a connected component (CC) of the graph, and those with between 18 and 24 nt are classified as putative miRNAs, while longer re-assembled unipaths (>24 nt) are classified as other longer sequences (Figure 1.6). BrumiR then restores missing connections by re-clustering the putative miRNAs performing an all-vs-all comparison. The most expressed miRNA is selected as the representative of the cluster (Figure 1.7) and the remaining members are classified as potential isomiRs (Figure 1.7). The final BrumiR step uses the RFAM database to discard predicted miRNAs matching to other classes of RNA (*e.g.* Ribosomal genes, Figure 1.8). As an example, BrumiR reduces the input sRNA-seq data by five orders of magnitude generating less than 1,000 putative mature miRNAs (24 million input reads to 966 miRNA candidates, see Figure 1.10). Finally, BrumiR outputs several FASTA files with all predicted mature miRNAs, all longer RNAs, putative isomiRs, other sRNAs (RFAM comparison), and a table with expression values for each predicted miRNA. Additionally, BrumiR outputs the final graph in GFA format, which can be explored using Bandage (43) (Figure S7).

**Figure 1.**
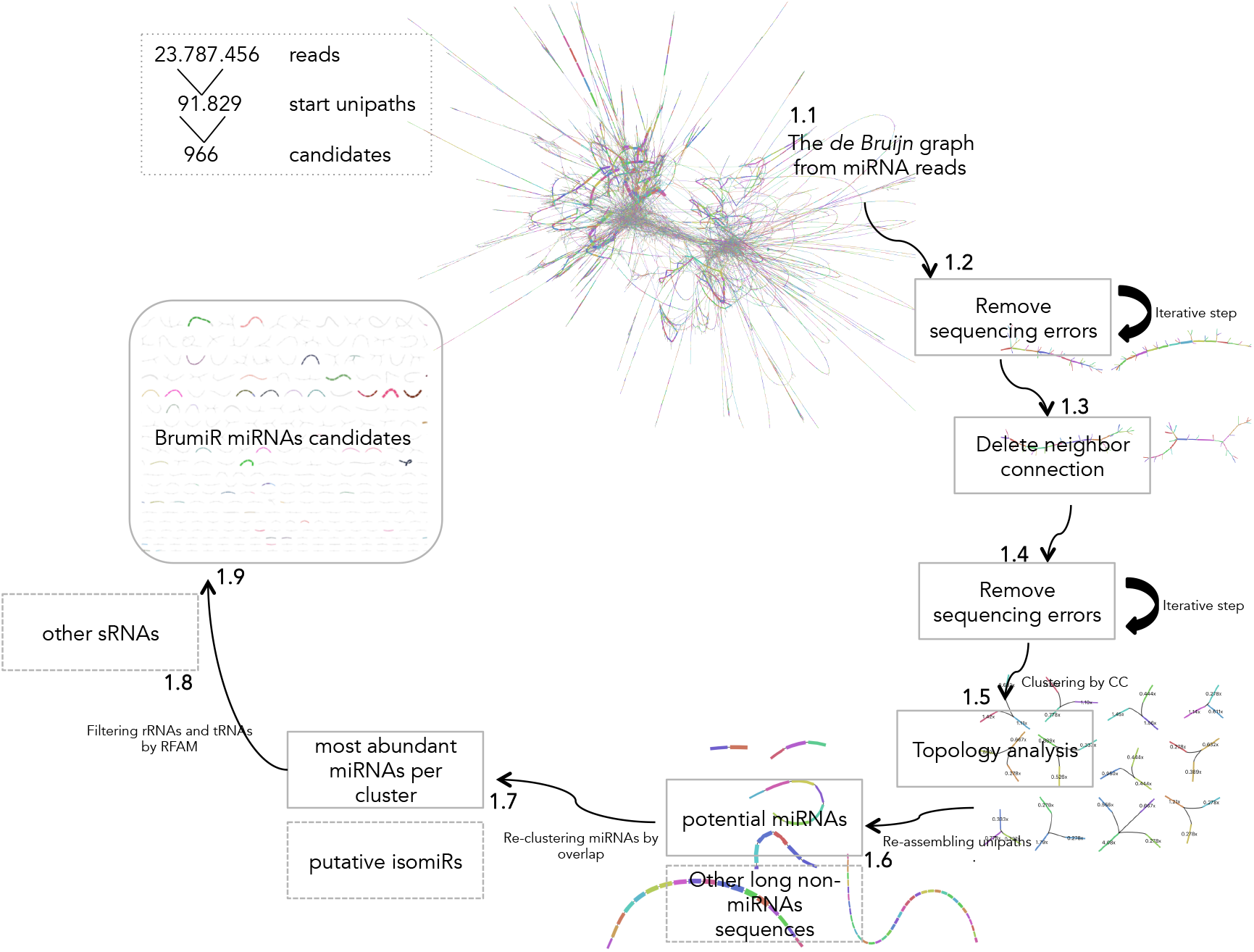
BrumiR algorithm. Different steps of BrumiR to discover miRNAs from sRNA-seq data. **1.1** De Bruijn graph step, **1.2** Tips removal iterative step, **1.3** Delete neigbor connection step, **1.4** Tips removal step repetition, **1.5** Topology analysis step, **1.6** Re-assembling unipaths by CC step, **1.7** Re-clustering by overlap step, **1.8** Filtering other sRNAs by RFAM step, **1.9** BrumiR candidates catalog.

### BrumiR achieves the highest accuracy on simulated data

To evaluate the performance of BrumiR, we applied it to discover mature miRNAs on simulated sRNA-seq reads from 10 animal and 10 plant species (Figure 2A). We compared BrumiR to the state-of-the-art genome-based miRNA discovery tools miRDeep2 (21) and miR-PREFeR (22), which were developed specifically for animal (miRDeep2) and plant (miR-PREFeR) species. For each tested species, we generated two synthetic datasets with different error-rates (0.01 and 0.02) using the miRsim tool implemented and provided by the BrumiR toolkit (https://github.com/camoragaq/miRsim), and the high-confidence miRNAs annotated in the miRBase database (see Methods section). A total of 20 datasets with an average of 11.5 million reads were simulated. The list of simulated miRNAs was considered as the ground truth, and benchmark metrics (Figure 2C) were computed to assess the performance of BrumiR and of the other software (See Methods section) (Supplementary Table S2).

**Figure 2.**
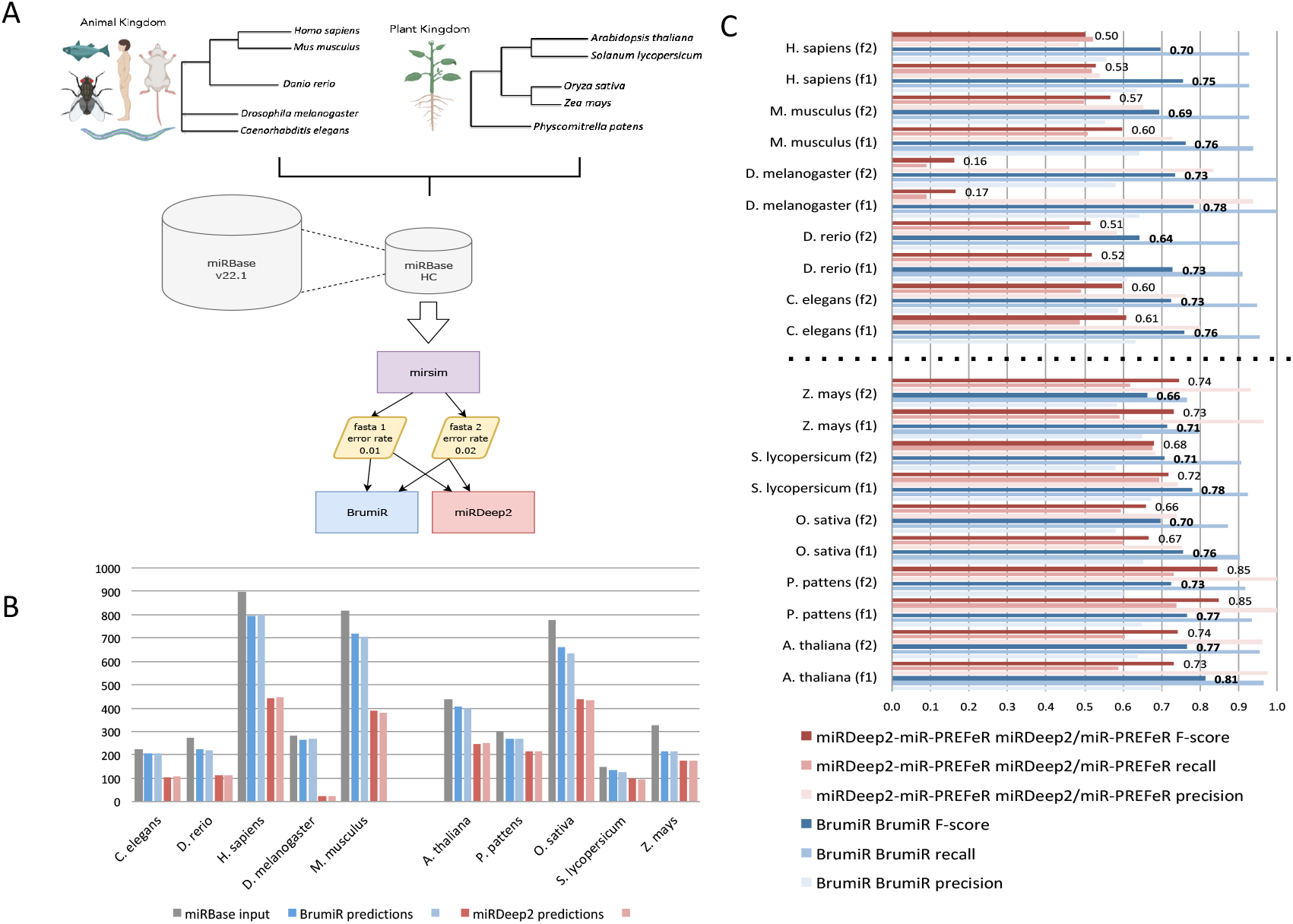
Synthetic benchmarking between BrumiR and miRDeep2. **A)** Workflow and species selected, **B)** miRBase input vs miRNA true positive predictions for each tool (2 samples), **C)** Benchmarking metrics for all datasets tested.

BrumiR recovered more mature miRNAs than the others, on average 92% (opposed to 41% and 64% for miRDeep2 and miR-PREFeR, respectively), and presented the highest average recall across all the simulated datasets (Figure 2B). BrumiR recovered more than 90% of the simulated mature miRNAs in 17 of the 20 simulated datasets (Figure 2B). In particular in the *H. sapiens*, *M. musculus* and *D. melanogaster* datasets, BrumiR recovered three times more candidates than MiRDeep2 (Figure 2B). As concerns precision, BrumiR tended to generate more putative candidates than MiR-PREFeR (median 316 vs 264) and miRDeep2 (median 445 vs 308). The slightly higher number of BrumiR candidates resulted in lower average precision than miRDeep2 (0.59 vs 0.69) and MiR-PREFeR (0.63 vs 0.87). This is due to the fact that BrumiR does not use the hairpin structure filter employed by the other software. If we consider both precision and recall (F-Score), BrumiR was the top performer in 16 of the 20 datasets evaluated (Figure 2C). With animal species, BrumiR always reached a higher F-score than miRDeep2. With plant species, BrumiR was better or comparable to miR-PREFeR on most datasets, except for *Z. mays* and *P. pattens* where miR-PREFeR reached a higher F-Score (Figure 2C).

In terms of computational time, BrumiR was the fastest method. In particular, BrumiR core was on average 30X faster than miRDeep2 and 10X times faster than MiR-PREFeR (see Table S3). The speed of BrumiR relies on efficient alignment-free and graph-based approaches.

Overall, we demonstrated with simulated data that BrumiR discovers putative mature miRNAs without a reference genome across different eukaryotic species achieving the highest accuracy and computational efficiency.

### The hairpin structure of mature miRNAs is found in most of the BrumiR candidates

In order to assess the performance of BrumiR on real data, we collected public datasets for the same plant and animal species evaluated in the synthetic benchmark (Figure 2A). On average, 15.4 and 18.2 raw million reads were used for the animal and plant datasets (Supplementary Table S2), respectively. The predictions of BrumiR were compared against those of the state-of-the-art tools encompassing reference and *de novo* based methods (14,21,22). In particular, we included mirnovo that similarly to BrumiR can discover mature miRNAs directly from the reads. Before running the tools, low-quality reads were removed using fastp (26) (~10%, see Methods section). All the predicted miRNAs for each tool were annotated using the miRBase database to identify known and novel predictions. On average, BrumiR predicted ~1,000 putative mature miRNAs for the animal species, which was ~2.7X higher than the miRDeep2 candidates and 1.7X lower than the candidates predicted by mirnovo (Figure 3A1). For plant species, BrumiR predicted on average ~1,900 putative mature miRNAs, which was lower than the candidates predicted by mirR-PREFeR (3,248 on average), and higher than the predictions of mirnovo (301 on average) (Figure 3A1). A comparison using the miRBase (13) annotated miRNAs revealed that BrumiR shared more candidates with miRDeep2 and miR-PREFeR than with mirnovo (Figure 3A2). However, an important fraction (on average more than 70%) of the miRBase-annotated candidates were exclusive to each tool (Figure 3A2), which summarizes the complexity of miRNA discovery.

**Figure 3.**
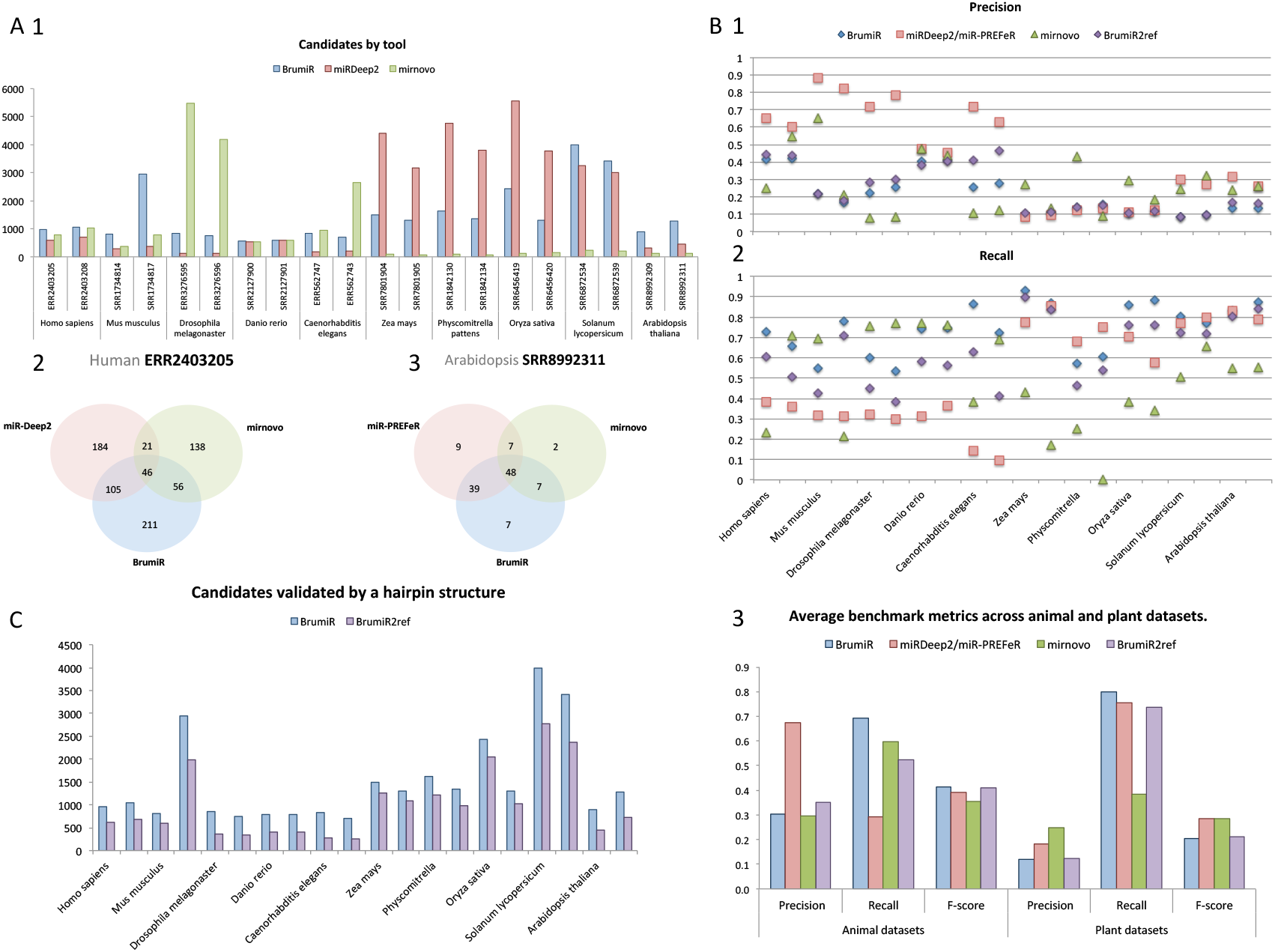
Real dataset benchmark of BrumiR and state-of-the-art tools. **A)** Number of predictions by tool for all the datasets and the overlap between them for 2 datasets (1 for animal and 1 for plant); **B)** Benchmarking metrics computed using miRBase annotated miRNAs, precision and recall for each dataset; and average metrics, including F-score. **C)** BrumiR candidates validated by Hairpin structure (BrumiR2Reference).

Considering miRBase-annotated candidates as the ground-truth, we computed precision, recall and F-Score for all the evaluated tools (Figure 3B, Method section). BrumiR achieved an accuracy (F-Score) better (animals) or comparable (plants) to the one obtained by the other software (Figure 3B3). Moreover, BrumiR consistently reached the highest recall for most of the datasets evaluated (Figure 3B2). The precision values of BrumiR were slightly lower for some datasets (Figure 3B1), however, none of the methods performed well on this metric (Figure 3B3). In particular, miRdeep2 reached the highest precision (~0.7) on animal species and all methods performed poorly on the evaluated plant species (average precision <0.3). The low precision with plant species may be the product of a low number of entries annotated in miRBase for plants (10.414 vs 38.471 animals) as well as of a higher complexity of plant miRNAs (38).

The BrumiR toolkit also provides a tool to determine the hairpin loop of miRNA precursor sequences, which is the main structural feature of miRNAs (40). BrumiR2reference maps the BrumiR predicted mature miRNA to the reference genome using an exhaustive alignment (See Methods section), generates precursor sequences, computes its secondary structure, and checks the hairpin structure using a variety of criteria inferred from analyzing more than 30,000 miRBase precursor sequences from animal and plant species (see Methods section). We used BrumiR2reference as a double validation for all the predicted mature miRNAs generated by BrumiR for the animal and plant datasets (Figure 3C). On average, BrumiR2reference identified a valid precursor sequence having the characteristic hairpin structure for over 70% of the BrumiR candidates (Figure 3C).

In terms of speed, BrumiR core was the fastest tool. BrumiR was on average 120X and 220X times faster than miRDeep2 and miR-PREFeR, respectively (See Table S4).

Overall, we demonstrated that BrumiR is a competitive tool for discovering mature miRNAs without a reference genome. We showed that it was the most sensitive on most of the datasets tested. The performance of our method was not only faster, but also better or comparable to the state-of-the-art tools. Moreover, we also provide a new mapper approach to be used when a reference genome is available, to further verify if a precursor sequence of the predicted mature miRNA is present in the genome. BrumiR therefore represents a reliable alternative for the discovery of mature miRNAs in model and non-model species with or without a reference genome.

### Discovering novel miRNAs from sRNA-seq data of *A. thaliana* roots using BrumiR

*A. thaliana* is one of the best characterized model organisms, and the first plant species in which miRNAs were cloned and sequenced (44). To date, 436 mature miRNA sequences are included in the miRBase database. Most of these miRNAs have been identified by studies addressing the sRNAome of different plant organs (45), cell types (46), or responses to biotic or abiotic stress using sRNA-seq (47) (48).

We sequenced sRNA-seq libraries from the roots of *A. thaliana* after different time points during vegetative development (see Methods section) (Figure S8) to demonstrate the potential of BrumiR to discover novel mature miRNAs in a known biological context. BrumiR was run independently for each condition and replicate. The day 5 samples were excluded because of the low number of reads when compared to the other samples (Table S5). BrumiR predicted, on average, 1,120 mature miRNAs per sample, which were further refined to 678 using the BrumiR2ref tool. To take advantage of our experimental design, we considered as a putative miRNA the ones present in the three replicates (core predictions) (49) (Figure 4A). Novel miRNAs were identified using the following steps: First, predictions were classified as known miRNAs by comparing with miRBase (160 known miRNAs out of a total of 436 miRNAs already described for *A. thaliana* in miRBase). These known miRNAs were put aside to explore the sensitivity of BrumiR in detecting novel putative miRNAs. We then clustered the remaining putative miRNAs into three stages: early, late, and constitutive (Figure 4B). The days 9, 13 and 17 represent an early stage of the plant development (50); days 17, 21 and 25 represent a late stage of the plant development (50), and the putative miRNAs expressed in all conditions represent the constitutive category. A total of 25 putative novel miRNAs were identified, and a manual curation was carried out using all the information provided by BrumiR.

**Figure 4.**
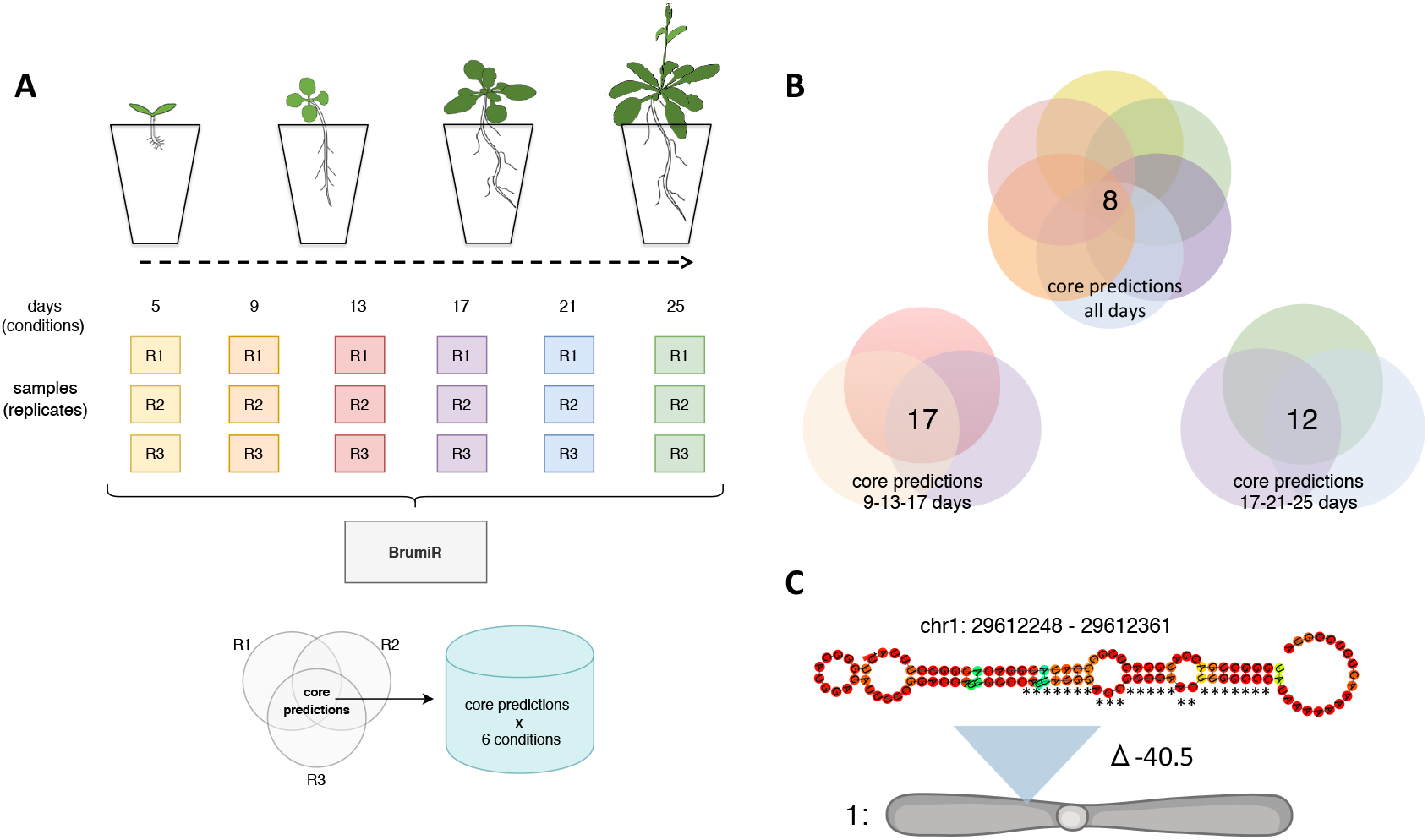
Applying BrumiR on sRNA-seq from *Arabidopsis* root libraries. **A)** Experimental design implemented; roots from *Arabidopsis* in a time-scale per day as conditions were sequenced in three technical replicates. BrumiR was used to analyze all sRNA-seq librairies, and conserved predictions by the three replicates was considered as a core by condition. **B)** Different combinations of root growth per day were analyzed together to identify novel putative miRNAs conserved in all conditions. **C)** miR-8 is discovered as a novel miRNA, supported by the hairpin analysis, and is conserved in all replicates in all conditions.

Three curated novel miRNAs fulfilling all the recommended criteria to annotate miRNAs in plants (49) were discovered by BrumiR (Figure S9) (Table S6). One of the curated miRNAs is located in chr1:29612248-29612361 (from now on denoted as miR-8) with a free energy of −40.5 and the characteristic hairpin structure of plant miRNAs (Figure 4C). Interestingly, this miRNA locus has not been previously discovered because its mature sequence maps to multiple chromosomes, and is therefore discarded by genome-based tools (17).

In an exploratory analysis to shed light on the potential targets of these novel miRNAs, we conducted an *in silico* target transcript prediction using the psRNATarget algorithm (42) (Supplementary Table S7). FSD-1 (AT4G25100) was found to be one of the top genes regulated by this novel miRNA miR-8 (Supplementary Table S6). In *A. thaliana*, FSD-1 encodes a Fe superoxide dismutase enzyme which regulates reactive oxygen species (ROS) levels of chloroplast and cytosol and participates in salt stress tolerance (51). Moreover, knockout mutants of FSD-1 exibit a lower number of lateral roots, thereby suggesting an important role in root development (52). FSD-1 is developmentally regulated, abundantly expressed from the 3^rd^ day to the 13^th^ day but significantly decreased in the following days, and its differential accumulation between root zones is related to emerging patterns of lateral roots and hair formation from trichomes (53). Another predicted mRNA target of miR-8 is PER24 (AT2G39040), which is a peroxidase gene involved in the detoxification of ROS in the extracellular and Na+ homeostasis and which plays an important role in the resistance to salinity stress as does the FSD-1 gene (51). It is plausible to say that these novel miRNAs may be involved in the fine-tuning of lateral root growth in the early stages of development. These results highlight the value of the BrumiR toolkit for discovering novel miRNA candidates with functional impact on the organisms studied, even in the case where high quality genomes are available.

## DISCUSSION

In this paper, we introduced and benchmarked the BrumiR toolkit, which was designed for enabling the identification of mature miRNAs in model and non-model species with or without a reference genome, encompassing the plant and animal kingdoms. The BrumiR toolkit implements the following algorithms: 1) a new discovery miRNA tool (BrumiR-core), 2) a specific genome mapper (BrumiR2ref), and 3) an sRNA-seq read simulator (miRsim). We demonstrated that BrumiR is capable of identifying mature miRNAs based only on the sequence information, and generates results that are better or comparable to the state-of-the-art tools on simulated and real datasets. We further tested the usefulness of the BrumiR toolkit for discovering novel miRNAs potentially involved in the regulation of the root development of the extensively annotated *A. thaliana* genome.

Unlike the state-of-the-art tools, BrumiR starts by encoding the sRNA-seq reads using a de Bruijn graph. This avoids the read mapping stage and the dependency on previous miRNA annotations. It also enables the identification of sequencing artifacts. A critical step of genome-based miRNA discovery tools is to identify the precursor sequence when a reference genome is available. BrumiR introduces a new mapping approach, BrumiR2ref, which scans every possible hairpin precursor in the genome, when such is available, for all the BrumiR predictions. As the hairpin structure is determined using the predicted mature miRNA instead of the reads, this alignment can support mismatches and indels and handles the case of multi-mapped candidates (due to repetitive regions of the genome). Such features distinguish BrumiR from the current genome-based methods.

Discovering miRNAs in non-model species is one of the limitations of the current methods. One exception is mirnovo, which similarly to BrumiR can predict miRNAs using only the sRNA-seq data, and a specific training set for animal and plant species. We thus compared its performance to the one of BrumiR. Our results show that mirnovo is very conservative, generating few predictions but with a high accuracy. BrumiR generates a larger number of candidates than mirnovo, most of which are annotated in miRBase but also others that correspond to new candidates. However, the higher number of predictions of BrumiR results in a lower accuracy in some of the datasets evaluated, representing a potential weakness of our method. To address this, we developed the BrumiR2ref tool to refine the BrumiR predictions, thus reducing the number of false-positive putative miRNAs. We plan to further reduce the false-positive rate of BrumiR by using a random-forest classifier trained on the high confidence mature sequences available in miRBase. The latter will improve the accuracy of BrumiR even in the case when a reference genome is not available. It is important to observe that the miRNA annotations remain incomplete and although miRBase is the main repository for miRNAs, it cannot be considered as the gold-standard for most species (many of the entries have not been correctly validated, for example) (9). For this reason, the predictions of BrumiR are not based on miRBase in any step of the algorithm. However, miRBase can be used with all due precaution in a posterior analysis to verify the miRNAs inferred in case of not having any reference genome. In terms of computational resources and usability, BrumiR is the fastest method and provides a stand-alone package for running locally all the analyses. It further generates an output that is compatible with the bandage software (43), which can be employed to visualize and explore the results of BrumiR in a user-friendly way.

Moreover, BrumiR reports other sequences expressed in the sRNA-seq data among which are putative isomiRs and longer non-coding RNAs, thereby providing additional biological insight.

Finally, we tested the effectiveness of BrumiR on sequenced sRNA-seq libraries from the roots of *A. thaliana*, and were able to annotate 3 novel putative miRNAs based on the very conservative criteria proposed in (49), showing the potential of it being used alone or in combination with other methods.

In summary, we present a new and versatile method that implements novel algorithmic ideas for the study of miRNAs that complements and extends the currently existing approaches.

## MATERIALS AND METHODS

### Building a de Bruijn graph for sRNA-seq data

BrumiR starts by building a compact de Bruijn graph from the sRNA-seq reads given as input. De Bruijn graphs are a widely used approach in the genome assembly problem (25). BrumiR uses this graph to organize, detect, and exploit the sequence information of sRNA-seq experiments. BrumiR takes as input sequencing files in FASTA or FASTQ formats. The sequencing data can be cleaned, using a fastq pre-processor (26) (*i.e.* fastp), to remove adapter sequences and trim low quality bases. BrumiR employs the BCALM (27) tool to build a de Bruijn graph from the sRNA-seq reads. BCALM uses a node-centric bi-directed de Bruijn graph where the nodes are *k*-mers, that is words of length *k*, and an arc between two nodes if the *k-1* suffix of one node is equal to the *k-1* prefix of the subsequent node, representing an exact overlap of *k-1* bases (27). A critical parameter of any de Bruijn graph approach is the *k*-mer size (28). We observed that the length of all mature miRNA sequences stored in the miRBase database (v21) (13) have a minimum value of 18 nt (Supplementary Figure S1). We thus empirically set the *k*-mer size equal to 18. BCALM compacts the nodes of the de Bruijn graph into maximal unipaths by gluing all the nodes of the graph with an in-degree and an out-degree equal to one, thus generating the so-called *unipath graph* (27). The unipath graph is the starting point of BrumiR (Figure1A). Notice that the unipath graph generated by BCALM does not represent what is expected for a set of mature miRNAs (one connected component for each miRNA) and therefore further graph operations are needed. BrumiR uses a minimum *k*-mer frequency (KM value) of 5 and all *k*-mers with lower frequency are ignored, without losing most of the information contained in the sequencing reads (Supplementary Figure S2).

### Removing sequencing errors from the unipath sRNA-seq graph

BrumiR deletes from the unipath graph all the nodes that have only one connection (degree equal to 1), known as dead-end paths or tips (30). Usually, these nodes have a low abundance value associated to them (KM less than or equal to 5, the default parameter). Moreover, BrumiR deletes isolated nodes (degree equal to 0) having a low abundance; isolated nodes highly expressed are however conserved for further analysis. All these nodes are likely artifacts generated from sequencing errors because they are not deeply expressed in the sRNAs-seq reads (29). BrumiR iterates this step 3 times in order to prune and clean the unipath graph (polishing). This operation, called ‘tip removal’, edits the original unipath graph, and therefore a new unipath graph with a new structure is generated (Figure 1B).

### An expressed mature miRNA has uniform coverage

The unipath graph of a set of miRNAs from an sRNA-seq experiment has non-uniform coverage as different miRNAs and other elements may be connected in a single big component (Figure 1.1). BrumiR evaluates each connection of the unipath graph to identify those that link two nodes with a large expression difference. According to the miRNA biogenesis, after a stable miRNA precursor is cleaved by Dicer, among its three products, the miRNA mature sequence is the most abundant and when it is sequenced, it has a uniform expression along its sequence (20). Thus due to miRNA biogenesis, it is possible to capture the complete miRNA mature sequence having a homogeneous expression (9) directly from the sRNA-seq experiments. BrumiR expects a similar KM value for *k*-mers originating from the same mature miRNA gene. Accordingly, if we observe two connected nodes that show a big difference in their abundance values, this connection is deleted and we keep the nodes unconnected. In particular, two unipaths *U*=*{a,b}* connected in the graph have a KM value associated to them that represents their coverage from the reads information. BrumiR scans all the neighbor connections and if the difference between their KMs is larger than three-fold, the connection is deleted (Ui_km_/Uj_km_ > 3). In this way, BrumiR defines a relative threshold that will depend on each unipath neighborhood in the graph. Finally, BrumiR repeats the tips removal step to eliminate new low frequency isolated nodes (Figure 1C).

### miRNAs and other sequences are captured in single connected components

After the previous steps of BrumiR, a new unipath graph emerges, with a new structure. It is thus necessary to identify and classify the new connected elements within the graph (Figure 1). A connected component (CC) of a graph is a maximal strongly connected subgraph (31). BrumiR computes the CCs of the unipath graph, and then each CC is processed independently to identify miRNA candidates as well as to discard other sequences present in the unipath graph.

### BrumiR classifies low abundance non-linear topologies as sequencing artefacts

BrumiR detects topologies that are potentially related to sequencing errors and thus unlikely to be miRNA candidates. The shapes of these topologies were identified by visual inspection of several unipath graphs and are described in detail in Figure S3. Usually they have low KM and are composed of lowly expressed branching nodes with 3, 4 or 5 connections to the principal structures in the graph (Figure S3). Moreover, we observed that the sequences contained in these topologies were usually redundant and contained in other linear and more expressed CCs. In this way, we are not discarding relevant sequence information. BrumiR removes about 10% of the CCs in this step.

### Re-assembling unipaths within each CC

BrumiR re-assembles all unipaths present in the linear CCs by bundling the nodes with in and out degree equal to 1 into a new unipath. BrumiR classifies them into different types based on their length. The latter is the length of the sequence represented by the new unipath. All CCs having a length between 18 and 24 are stored as potential miRNA sequences. The CCs corresponding to an isolated node that have high KM (KM>50) are included in the latter group. CCs with lengths over 24 are classified as longer sequences or other types of genomic sequences captured along with the miRNAs. The longer sequences are put aside for later analysis. Moreover, BrumiR identifies circular CCs and branching CCs. The former are circular unipaths and the latter CCs with a high number of branching nodes. Branching CCs are not considered in the subsequent steps because they are likely sequencing errors (low abundance) or contamination present in the sRNA-seq data (Figure S4).

### Re-clustering potential miRNAs

After grouping unipaths by CCs, BrumiR builds an overlap graph to rescue the missing connections between potential miRNA candidates sharing an overlap with another candidate. First, BrumiR adds all the candidates as nodes of the overlap graph, then an all-vs-all *k*-mer comparison is performed using exact overlaps of length *k*=15. Candidates sharing an exact overlap are connected in the overlap graph. Then, the connected components are computed to identify clusters of miRNA candidates, and the most expressed candidate within each component is selected as the representative candidate of the cluster. The representative candidates are compared all-vs-all in a second overlap step that allows a maximum edit distance of 2, which is implemented using the edlib library (32). BrumiR then builds a second overlap graph, computes again the connected components, and selects the most expressed candidate as the representative of each cluster. The other members of each connected component are classified as putative isomiRs and saved in a file for later analysis.

### Identifying other expressed RNA sequences

In sRNA-seq experiments, different types of RNAs are expressed, some of which, such as small non-coding RNA elements, may have similar length with miRNAs (33). The RFAM database (34) is a collection of curated RNA families including three functional classes of RNAs (non-coding, cis-regulatory elements, and self-splicing RNAs), which are classified into families according to their secondary structure and sequence information (Covariance Models) (34). We downloaded 3,017 RNA families present in RFAM (v14.1) and excluded 529 miRNA families (35). The sequences of 2,488 RFAM families were concatenated (a total of 2,736,549 sequences) and used to build a 16-mer database with the KMC3 *k*-mer counter tool (36) (“-fm -n100 -k16 -ci5”). All distinct 16-mers with a frequency lower than 5 were excluded, leading to a total of 6,204,556 distinct 16-mers related to RNA elements. Additionally, we downloaded all the mature miRNA sequences from miRBase (v22.1) (13) and built a 16-mer database with KMC3 (“-fm -n100 -k16 -ci1 mature.fa.gz”). RFAM 16-mers matching 16-mers from the 16-mer mature miRBase database were excluded from RFAM, leading to a 16-mer RFAM database with a total of 6,204,487 distinct 16-mers. Finally, the BrumiR candidates (18-24 length) were matched to the 16-mer RFAM database, and matching candidates were excluded and reported as sequences potentially associated to other RNA elements. The BrumiR candidates passing the aforementioned filter are reported as the final list of miRNA candidates.

### Identifying precursor sequences for BrumiR candidates (BrumiR2Reference)

Unlike current state-of-the-art tools that perform miRNA discovery by mapping the sRNA-seq reads to a reference genome, BrumiR generates candidates by operating directly on the sRNA-seq reads. The reduced list of potential BrumiR miRNA candidates permits the computation of a more exhaustive alignment than when mapping directly the sRNA-seq reads to the reference genome. BrumiR aligns each candidate to the reference genome using an exact alignment method that computes the edit distance (38) between two strings and thus support mismatches, insertions and deletions. The BrumiR2reference tool divides the reference genome in non-overlapping windows of 200bp (adjustable parameter), then the window is indexed using 12-mers and each miRNA candidate is matched in both strands (split at 12-mers). When a 12-mer match is found, an exhaustive alignment is computed between the window and the matching miRNA candidate. The alignment is performed using a fast implementation of Myers’ bit-vector algorithm (32). A miRNA candidate is stored as hit if the alignment in the current genomic window has an edit distance less than or equal to 2. After scanning all the genomic windows, the vector of hits is sorted by miRNA-candidate; edit distance (0-2), and alignment sequence coverage. For a single miRNA-candidate, a maximum of 100 genomic locations (best hits) are selected. BrumiR2reference then builds a potential precursor sequence for each selected hit using a strategy similar to the ones employed by miRDeep2 (21) and Mirinho (37). BrumiR excises the potential precursor hairpin sequence from the flanking genomic coordinates of the reported miRNA candidate hits (mature sequence) in both strands. Potential precursor hairpin sequences of length 110 bp are built for animal species from both strands, while for plant species hairpin sequences of lengths 110, 150, 200, 250 and 300 bp are built from both strands (38). Secondary structure prediction for all the potential precursor sequences is performed using RNAfold (v2.4.9) (39). Secondary structures with a minimum free energy in the range of 15-80 kcal/mol are checked for a hairpin loop characteristic of miRNAs (40) (Figure S5). Structures with a hairpin loop composed of a single segment without pseudo-knot, multi-loops, external loops and with less than 5 bulges, 3 dangling ends, and 10 internal loops are classified as characteristic secondary structures of miRNA precursor sequences. The aforementioned filters were derived from analyzing the secondary structure of 38,589 precursor sequences stored in miRBase (v22.1) (13) using a modified version of the bpRNA program (41) (Figure S6).

### Benchmarking BrumiR using simulated sRNA-seq reads

We simulated synthetic reads from animal and plant species, and compared the results of BrumiR to those obtained with the miRDeep2 (21) and miR-PREFeR (22) tools. The sRNA-seq reads were simulated using miRsim (https://github.com/camoragaq/miRsim), a tool that we developed specifically for simulating sRNA-seq reads from a list of known miRNA mature sequences. miRsim is based on *wgsim* (https://github.com/lh3/wgsim), which is a widely used tool for simulating short Illumina genomic reads. miRsim includes functionalities specific of sRNA-seq reads such as variable depth/coverage and shorter read lengths. miRNA mature sequences were obtained from miRBase (13) for animal (High Confidence) and plant species. The animal species that we considered were: *Homo sapiens, Mus musculus*, *Drosophila melanogaster, Danio rerio*, and *Caenorhabditis elegans*, while the following plant species were included: *A. thaliana*, *Oryza sativa, Physcomitrella patens, Zea mays,* and *Solanum lycopersicum*. Supplementary Table S1 provides further details (*i.e.* number of reads, number of mature miRNAs *etc*.) for each simulated dataset. MiRDeep2 was run on the animal datasets with the default parameters providing the respective reference genome. Similarly, miR-PREFeR was run with the default parameters on the plant datasets. BrumiR was run with the default parameters on both the animal and plant datasets. The list of simulated miRNAs was considered as the ground truth, and precision, recall and F-Score quality metrics were computed to assess the performance of each discovery tool. The benchmark metrics were defined as follows:

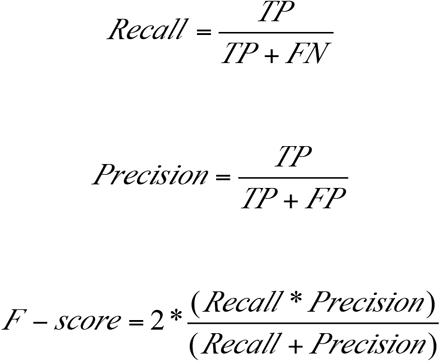

where:

TP= true positive elements predicted as miRNAs present in the miRBase input list.
FP= false positive elements predicted as miRNAs but not present in the miRBase input list.
FN= false negative elements not predicted as miRNAs, but that were present in the miRBase input list.

### Benchmarking BrumiR using real sRNA-seq reads

We downloaded publicly available sRNA-seq data for the plant and animal species listed in the synthetic benchmark, and two datasets for each species were included (Supplementary Table S2). Additionally, we included mirnovo (14), a tool that can discover miRNAs without a reference genome. The predictions of BrumiR were benchmarked along with MiRDeep2 (v2.0.1.2) (21) and mirnovo for the animal datasets. Similarly, miR-PREFeR (22) replaced MiRDeep2 for the plant datasets (Supplementary Table S2).

The stand-alone packages of BrumiR, miRDeep2 and miR-PREFeR were used to discover miRNAs in all datasets. The software mirnovo was run using its web version because the stand-alone package was not available and the developer recommends the use of the web version instead. The miRNA discovery was performed for each sample independently using default parameters for MiRDeep2, miR-PREFeR and mirnovo. In particular, we used the scripts provided by miRDeep2 and miR-PREFeR to map the reads to the reference genome, and the predictions for these tools were performed on the resulting alignment files. The mirnovo predictions were done using the animal and plant universal panel respectively, as recommended when the reference genome is not available. BrumiR was run using the command line and parameters provided in the Supplementary Section 1. Moreover, the predictions of BrumiR were refined using the BrumiR2reference tool on the available reference genome of the selected species (Supplementary Table S2). Benchmark metrics (precision, recall, and F-Score) were computed as before but considering all the annotated mature sequences present in miRBase (v22.1) (13) as the ground-truth.

### miRNA discovery from *Arabidopsis* root samples

*A. thaliana* Col-0 seedlings were grown hydroponically on Phytatrays on 0.5X Murashige and Skoog medium (Phytotechnology Laboratories, cat. M519) under long-day conditions (16h light and 8h dark) at 22°C. Total RNA was isolated from plant roots after 5, 9, 13, 17, 21, and 25 days post-germination using the mirVana miRNA Isolation Kit (Thermo Fisher Scientific, cat. AM1560). RNA concentration was determined using the Qubit RNA BR Assay Kit (Thermo Fisher Scientific, cat. Q10210), and integrity was verified by capillary electrophoresis on a Fragment Analyzer^TM^ (Advanced Analytical Technologies, Inc.). The indexed sRNA libraries were built employing the TruSeq small RNA Sample Preparation Kit (Illumina, Inc.) following the manufacturer’s instructions. Briefly, 3’ and 5’ adaptors were sequentially ligated to 1 μg of total RNA prior to reverse transcription and library amplification by PCR. Size selection of the sRNA libraries was performed on 6% Novex TBE PAGE Gels (Thermo Fisher Scientific, cat. EC6265BOX) and purified by ethanol precipitation. Both the library size assessment and library quantification were carried out in a Fragment Analyzer^TM^. Finally, the libraries were pooled and sequenced on an Illumina NextSeq 500 platform.

All samples were analyzed with BrumiR separately with default parameters to identify the candidate miRNAs. We further validated the candidates having a putative precursor with a hairpin structure analysis using the BrumiR2ref tool with the reference genome for *A. thaliana* (GCF_000001735.4_TAIR10.1_genomic.fna). All validated candidate miRNAs were compared with known miRNAs described for *A. thaliana* (437) present in miRBase (v21). The putative novel miRNAs were curated manually, specifically, we checked the hairpin features, mature sequence alignment position, star sequence in the precursor sequence, mismatches in the seed region, and exact overlap in the antisense miRNA sequence (9). Then a target analysis was performed using the Araport 11 cDNA library with the plant-specific psRNATarget algorithm (based on a best expectation score) (42).

## Supporting information

Supplementary files of BrumiR

## CODE AVAILABILITY

The BrumiR code (v1.0) used in this manuscript is freely available at https://github.com/camoragaq/BrumiR, and is open software under the MIT license.

## ACKNOWLEDGEMENTS

This work was supported by CONICYT BECAS CHILE DOCTORADO 2016/FOLIO 72170320 granted to CM, by a post-doctorate fellowship from the Agence National de Recherche (ANR-GREEN 17_CE20_0031_01) granted to MGF, as well as by Fondo Nacional de Desarrollo Científico y Tecnológico (FONDECYT)-ANID grant 1170926, ANID PCI-Redes Internacionales entre Centros de Investigación grant REDES180097 and Instituto Milenio iBio - Iniciativa Científica Milenio MINECON granted to E.A.V. This research was performed using the computing facilities of the LBBE/PRABI and the France Génomique e-infrastructure (ANR-10-INBS-09-08).

## AUTHOR CONTRIBUTIONS

CM designed, developed, implemented and benchmarked BrumiR. MFS guided the development of BrumiR. ES conducted the *A. thaliana* experiments. EAV designed and supervised the *A. thaliana* experiments. CM wrote the initial version of the manuscript with inputs from all other authors. MFS and MGF helped to improve the manuscript. EAV, MFS, MGF and RAG provided crucial biological feedback. All authors provided helpful discussions for the work and reviewed the manuscript.

## References

1. Bartel DP. MicroRNAs: Genomics, Biogenesis, Mechanism, and Function. Cell. 2004 Jan 23;116(2):281–97.

2. Bartel DP. MicroRNAs: Target Recognition and Regulatory Functions. Cell. 2009 Jan 23;136(2):215–33.

3. Peng Y, Croce CM. The role of MicroRNAs in human cancer. Signal Transduct Target Ther. 2016 Jan 28;1:15004.

4. Greene J, Baird A-M, Brady L, Lim M, Gray SG, McDermott R, et al. Circular RNAs: Biogenesis, Function and Role in Human Diseases. Front Mol Biosci. 2017;4:38.

5. Lin R, He L, He J, Qin P, Wang Y, Deng Q, et al. Comprehensive analysis of microRNA-Seq and target mRNAs of rice sheath blight pathogen provides new insights into pathogenic regulatory mechanisms. DNA Res. 2016 Oct 1;23(5):415–25.

6. Wang M, Weiberg A, Lin F-M, Thomma BPHJ, Huang H-D, Jin H. Bidirectional cross-kingdom RNAi and fungal uptake of external RNAs confer plant protection. Nature Plants. 2016 Oct;2(10):16151.

7. Lau NC, Lim LP, Weinstein EG, Bartel DP. An Abundant Class of Tiny RNAs with Probable Regulatory Roles in Caenorhabditis elegans. Science. 2001 Oct 26;294(5543):858–62.

8. Lagos-Quintana M, Rauhut R, Lendeckel W, Tuschl T. Identification of novel genes coding for small expressed RNAs. Science. 2001 Oct 26;294(5543):853–8.

9. Bortolomeazzi M, Gaffo E, Bortoluzzi S. A survey of software tools for microRNA discovery and characterization using RNA-seq. Brief Bioinformatics. 2019 21;20(3):918–30.

10. Pinzón N, Li B, Martinez L, Sergeeva A, Presumey J, Apparailly F, et al. microRNA target prediction programs predict many false positives. Genome Res. 2017 Feb 1;27(2):234–45.

11. Morin RD, O’Connor MD, Griffith M, Kuchenbauer F, Delaney A, Prabhu A-L, et al. Application of massively parallel sequencing to microRNA profiling and discovery in human embryonic stem cells. Genome Res. 2008 Apr;18(4):610–21.

12. Chen L, Heikkinen L, Wang C, Yang Y, Sun H, Wong G. Trends in the development of miRNA bioinformatics tools. Brief Bioinformatics. 2019 27;20(5):1836–52.

13. Kozomara A, Griffiths-Jones S. miRBase: annotating high confidence microRNAs using deep sequencing data. Nucleic Acids Res. 2014 Jan 1;42(Database issue):D68–73.

14. Vitsios DM, Kentepozidou E, Quintais L, Benito-Gutiérrez E, van Dongen S, Davis MP, et al. Mirnovo: genome-free prediction of microRNAs from small RNA sequencing data and single-cells using decision forests. Nucleic Acids Res. 2017 Dec 1;45(21):e177–e177.

15. Langmead B, Trapnell C, Pop M, Salzberg SL. Ultrafast and memory-efficient alignment of short DNA sequences to the human genome. Genome Biology. 2009 Mar 4;10(3):R25.

16. Li H, Durbin R. Fast and accurate short read alignment with Burrows–Wheeler transform. Bioinformatics. 2009 Jul 15;25(14):1754–60.

17. Ziemann M, Kaspi A, El-Osta A. Evaluation of microRNA alignment techniques. RNA. 2016 Aug 1;22(8):1120–38.

18. Li Y, Zhang Z, Liu F, Vongsangnak W, Jing Q, Shen B. Performance comparison and evaluation of software tools for microRNA deep-sequencing data analysis. Nucleic Acids Res. 2012 May;40(10):4298–305.

19. A reference standard for genome biology. Nature Biotechnology. 2018 Dec;36(12):1121.

20. Friedländer MR, Chen W, Adamidi C, Maaskola J, Einspanier R, Knespel S, et al. Discovering microRNAs from deep sequencing data using miRDeep. Nature Biotechnology. 2008 Apr;26(4):407–15.

21. Friedländer MR, Mackowiak SD, Li N, Chen W, Rajewsky N. miRDeep2 accurately identifies known and hundreds of novel microRNA genes in seven animal clades. Nucleic Acids Res. 2012 Jan 1;40(1):37–52.

22. Lei J, Sun Y. miR-PREFeR: an accurate, fast and easy-to-use plant miRNA prediction tool using small RNA-Seq data. Bioinformatics. 2014 Oct;30(19):2837–9.

23. Jha A, Shankar R. miReader: Discovering Novel miRNAs in Species without Sequenced Genome. PLOS ONE. 2013 Jun 21;8(6):e66857.

24. Mapleson D, Moxon S, Dalmay T, Moulton V. MirPlex: a tool for identifying miRNAs in high-throughput sRNA datasets without a genome. J Exp Zool B Mol Dev Evol. 2013 Jan;320(1):47–56.

25. Compeau PEC, Pevzner PA, Tesler G. Why are de Bruijn graphs useful for genome assembly? Nat Biotechnol. 2011 Nov 8;29(11):987–91.

26. Chen S, Zhou Y, Chen Y, Gu J. fastp: an ultra-fast all-in-one FASTQ preprocessor. Bioinformatics. 2018 Sep 1;34(17):i884–90.

27. Chikhi R, Limasset A, Medvedev P. Compacting de Bruijn graphs from sequencing data quickly and in low memory. Bioinformatics. 2016 Jun 15;32(12):i201–8.

28. Durai DA, Schulz MH. Informed kmer selection for de novo transcriptome assembly. Bioinformatics. 2016 Jun 1;32(11):1670–7.

29. Deorowicz S, Debudaj-Grabysz A, Grabowski S. Disk-based k-mer counting on a PC. BMC Bioinformatics. 2013 May 16;14(1):160.

30. Chikhi R, Rizk G. Space-efficient and exact de Bruijn graph representation based on a Bloom filter. Algorithms for Molecular Biology. 2013 Sep 16;8(1):22.

31. Lewis HR, Papadimitriou CH. Symmetric space-bounded computation. Theoretical Computer Science. 1982 Aug 1;19(2):161–87.

32. Šošić M, Šikić M. Edlib: a C/C ++ library for fast, exact sequence alignment using edit distance. Bioinformatics. 2017 May 1;33(9):1394–5.

33. Lambert M, Benmoussa A, Provost P. Small Non-Coding RNAs Derived from Eukaryotic Ribosomal RNA. Non-Coding RNA. 2019 Mar;5(1):16.

34. Kalvari I, Argasinska J, Quinones-Olvera N, Nawrocki EP, Rivas E, Eddy SR, et al. Rfam 13.0: shifting to a genome-centric resource for non-coding RNA families. Nucleic Acids Res. 2018 Jan 4;46(D1):D335–42.

35. Kalvari I, Nawrocki EP, Argasinska J, Quinones-Olvera N, Finn RD, Bateman A, et al. Non-Coding RNA Analysis Using the Rfam Database. Current Protocols in Bioinformatics. 2018;62(1):e51.

36. Kokot M, Dlugosz M, Deorowicz S. KMC 3: counting and manipulating k-mer statistics. Bioinformatics. 2017 Sep 1;33(17):2759–61.

37. Higashi S, Fournier C, Gautier C, Gaspin C, Sagot M-F. Mirinho: An efficient and general plant and animal pre-miRNA predictor for genomic and deep sequencing data. BMC Bioinformatics. 2015 May 29;16(1):179.

38. Meyers BC, Axtell MJ, Bartel B, Bartel DP, Baulcombe D, Bowman JL, et al. Criteria for Annotation of Plant MicroRNAs. The Plant Cell. 2008 Dec 1;20(12):3186–90.

39. Lorenz R, Bernhart SH, Höner zu Siederdissen C, Tafer H, Flamm C, Stadler PF, et al. ViennaRNA Package 2.0. Algorithms for Molecular Biology. 2011 Nov 24;6(1):26.

40. Roden C, Gaillard J, Kanoria S, Rennie W, Barish S, Cheng J, et al. Novel determinants of mammalian primary microRNA processing revealed by systematic evaluation of hairpin-containing transcripts and human genetic variation. Genome Res. 2017 Mar 1;27(3):374–84.

41. Danaee P, Rouches M, Wiley M, Deng D, Huang L, Hendrix D. bpRNA: large-scale automated annotation and analysis of RNA secondary structure. Nucleic Acids Res. 2018 Jun 20;46(11):5381–94.

42. Dai X, Zhuang Z, Zhao PX. psRNATarget: a plant small RNA target analysis server (2017 release). Nucleic Acids Res. 2018 Jul 2;46(W1):W49–54.

43. Wick RR, Schultz MB, Zobel J, Holt KE. Bandage: interactive visualization of de novo genome assemblies. Bioinformatics. 2015 Oct 15;31(20):3350–2.

44. Reinhart BJ, Weinstein EG, Rhoades MW, Bartel B, Bartel DP. MicroRNAs in plants. Genes Dev. 2002 Jul 1;16(13):1616–26.

45. Fahlgren N, Howell MD, Kasschau KD, Chapman EJ, Sullivan CM, Cumbie JS, et al. High-Throughput Sequencing of Arabidopsis microRNAs: Evidence for Frequent Birth and Death of MIRNA Genes. PLOS ONE. 2007 Feb 14;2(2):e219.

46. Breakfield NW, Corcoran DL, Petricka JJ, Shen J, Sae-Seaw J, Rubio-Somoza I, et al. High-resolution experimental and computational profiling of tissue-specific known and novel miRNAs in Arabidopsis. Genome Res [Internet]. 2011 Sep 22 [cited 2020 Jul 29]; Available from: http://genome.cshlp.org/content/early/2011/11/20/gr.123547.111

47. Hsieh L-C, Lin S-I, Shih AC-C, Chen J-W, Lin W-Y, Tseng C-Y, et al. Uncovering small RNA-mediated responses to phosphate deficiency in Arabidopsis by deep sequencing. Plant Physiol. 2009 Dec;151(4):2120–32.

48. Moldovan D, Spriggs A, Yang J, Pogson BJ, Dennis ES, Wilson IW. Hypoxia-responsive microRNAs and trans-acting small interfering RNAs in Arabidopsis. J Exp Bot. 2010 Jan;61(1):165–77.

49. Axtell MJ, Meyers BC. Revisiting Criteria for Plant MicroRNA Annotation in the Era of Big Data. The Plant Cell. 2018 Feb 1;30(2):272–84.

50. Satbhai SB, Ristova D, Busch W. Underground tuning: quantitative regulation of root growth. J Exp Bot. 2015 Feb;66(4):1099–112.

51. Guan Q, Wu J, Yue X, Zhang Y, Zhu J. A nuclear calcium-sensing pathway is critical for gene regulation and salt stress tolerance in Arabidopsis. PLoS Genet. 2013 Aug;9(8):e1003755.

52. Kuo WY, Huang CH, Liu AC, Cheng CP, Li SH, Chang WC, et al. CHAPERONIN 20 mediates iron superoxide dismutase (FeSOD) activity independent of its co-chaperonin role in Arabidopsis chloroplasts. New Phytol. 2013 Jan;197(1):99–110.

53. Dvořák P, Krasylenko Y, Ovečka M, Basheer J, Zapletalová V, Šamaj J, et al. FSD1: developmentally-regulated plastidial, nuclear and cytoplasmic enzyme with anti-oxidative and osmoprotective role. Plant Cell Environ. 2020 Apr 25;

