## Supplementary files of BrumiR for "BrumiR: A toolkit for *de novo* discovery of microRNAs from sRNA-seq data"

Table S7 Novel microRNAs and their putative interactions obtained using psRNATarget. 10

### BrumiR commands

#### BrumiR commands for real and simulated benchmark.

```
#trimming raw sequences using fastp
fastp --adapter_fasta ../adapters.fa -i <prefix>.fastq.gz -o <prefix>.trim.fastq.gz

#running miRDeep2
mapper.pl <prefix>.fa -c -p genome-index -m -q -s <prefix>.reads_collapsed.fa -t <prefix>.reads_collapsed_vs_genome.arf -v -o 2
miRDeep2.pl <prefix>.reads_collapsed.fa genome.fna <prefix>.reads_collapsed_vs_genome.arf none none none 2><prefix>.report.log

#running miR-PREFeR
python process-reads-fasta.py samplelist.txt <prefix>.fa <prefix>.fa
python bowtie-align-reads.py -p 2 -k 20 -f -r genome.fna <prefix>.fa.processed
python miR_PREFeR.py -L -k pipeline config.file

#running BrumiR
perl brumir.pl -a <prefix>.trim.fastq.gz -p <prefix> -T 10 -R 2 > <prefix>.log

#running BrumiR2reference
#for animal species
perl brumir2reference.pl -a <prefix>.candidate_miRNA.fasta -b genome.fna -t 4 -p <prefix>

#for plant species
perl brumir2reference.pl -a <prefix>.candidate_miRNA.fasta -b genome.fna -t 4 -p <prefix> -x 1
```

Table S1: Simulated sRNA-seq used to evaluate the performance of BRUMIR.

| | | #reads (M) | %mapped (ref. genome) | unipaths | BrumiR candidates | BrumiR TP | BrumiR hit miRBase | miRDeep2/miR-PREFeR candidates | miRDeep2/miR-PREFeR TP | miRDeep2/miR-PREFeR hit miRBase | BrumiR precision | BrumiR recall | BrumiR F-score | miRDeep2/miR-PREFeR precision | miRDeep2/miR-PREFeR recall | miRDeep2/miR-PREFeR F-score |
|---|---|---|---|---|---|---|---|---|---|---|---|---|---|---|---|---|
| Animal datasets | | | | | | | | | | | | | | | | |
| *Homo sapiens* | f1 | 22.91 | 91.71 | 79,986 | 1,205 | 766 | 794 | 1,136 | 614 | 443 | 0.64 | 0.93 | 0.75 | 0.54 | 0.52 | 0.53 |
| | f2 | 22.91 | 89.32 | 108,829 | 1,391 | 776 | 798 | 1,298 | 627 | 449 | 0.56 | 0.93 | 0.70 | 0.48 | 0.52 | 0.50 |
| *Mus musculus* | f1 | 21.29 | 93.49 | 73,003 | 1,117 | 716 | 719 | 700 | 509 | 389 | 0.64 | 0.94 | 0.76 | 0.73 | 0.51 | 0.60 |
| | f2 | 21.29 | 91.07 | 99,995 | 1,284 | 710 | 707 | 767 | 501 | 380 | 0.55 | 0.93 | 0.69 | 0.65 | 0.50 | 0.57 |
| *Drosophila melagonaster* | f1 | 7.46 | 95.56 | 26,785 | 418 | 269 | 265 | 32 | 30 | 24 | 0.64 | 1.00 | 0.78 | 0.94 | 0.09 | 0.17 |
| | f2 | 7.46 | 92.43 | 36,683 | 472 | 274 | 270 | 36 | 30 | 24 | 0.58 | 1.00 | 0.73 | 0.83 | 0.09 | 0.16 |
| *Danio rerio* | f1 | 7.00 | 91.02 | 24,073 | 337 | 205 | 224 | 306 | 182 | 113 | 0.61 | 0.91 | 0.73 | 0.59 | 0.46 | 0.52 |
| | f2 | 7.00 | 88.05 | 33,370 | 396 | 198 | 221 | 309 | 180 | 113 | 0.50 | 0.90 | 0.64 | 0.58 | 0.46 | 0.51 |
| *Caenorhabditis elegans* | f1 | 5.42 | 98.01 | 20,082 | 323 | 204 | 205 | 196 | 157 | 105 | 0.63 | 0.95 | 0.76 | 0.80 | 0.49 | 0.61 |
| | f2 | 5.42 | 94.62 | 28,449 | 351 | 206 | 205 | 206 | 157 | 106 | 0.59 | 0.95 | 0.73 | 0.76 | 0.49 | 0.60 |
| Plant datasets | | | | | | | | | | | | | | | | |
| *Zea mays* | f1 | 8.44 | 79.17 | 22,503 | 230 | 149 | 215 | 166 | 160 | 159 | 0.65 | 0.80 | 0.71 | 0.96 | 0.59 | 0.73 |
| | f2 | 8.44 | 76.72 | 29,232 | 252 | 147 | 215 | 188 | 175 | 174 | 0.58 | 0.77 | 0.66 | 0.93 | 0.62 | 0.74 |
| *Physcomitrella patens* | f1 | 7.84 | 83.82 | 22,247 | 299 | 194 | 270 | 184 | 184 | 213 | 0.65 | 0.93 | 0.77 | 1.00 | 0.74 | 0.85 |
| | f2 | 7.84 | 80.76 | 29,796 | 333 | 200 | 267 | 184 | 184 | 213 | 0.60 | 0.92 | 0.73 | 1.00 | 0.73 | 0.85 |
| *Oryza sativa* | f1 | 19.92 | 72.25 | 61,034 | 849 | 553 | 662 | 1,296 | 974 | 440 | 0.65 | 0.90 | 0.76 | 0.75 | 0.60 | 0.67 |
| | f2 | 19.92 | 69.76 | 83,602 | 955 | 554 | 634 | 1,408 | 1,038 | 433 | 0.58 | 0.87 | 0.70 | 0.74 | 0.59 | 0.66 |
| *Solanum lycopersicum* | f1 | 3.52 | 75.33 | 11,751 | 180 | 121 | 133 | 1,078 | 798 | 100 | 0.67 | 0.92 | 0.78 | 0.74 | 0.69 | 0.72 |
| | f2 | 3.52 | 73.42 | 16,041 | 209 | 121 | 126 | 979 | 669 | 94 | 0.58 | 0.91 | 0.71 | 0.68 | 0.68 | 0.68 |
| *Arabidopsis thaliana* | f1 | 11.22 | 83.15 | 32,464 | 510 | 358 | 405 | 249 | 243 | 246 | 0.70 | 0.96 | 0.81 | 0.98 | 0.59 | 0.73 |
| | f2 | 11.22 | 80.37 | 43,174 | 571 | 365 | 399 | 278 | 267 | 253 | 0.64 | 0.95 | 0.77 | 0.96 | 0.61 | 0.74 |

Table S2: Real sRNA-seq data used to evaluate the performance of BRUMIR.

| | | *Homo sapiens* | | *Mus* musculus | | *Drosophila* melagonaster | | *Danio* rerio | | *Caenorhabditis elegans* | | *Zea mays* | | *Physcomitrella patens* | | *Oryza sativa* | | Solanum *lycopersicum* | | Arabidopsis *thaliana* | |
|---|---|---|---|---|---|---|---|---|---|---|---|---|---|---|---|---|---|---|---|---|---|
| | | *ERR2403205* | *ERR2403208* | *SRR1734814* | *SRR1734817* | *ERR3276595* | *ERR3276596* | *SRR2127900* | *SRR2127901* | *ERR562747* | *ERR562743* | *SRR7801904* | *SRR7801905* | *SRR1842130* | *SRR1842134* | *SRR6456419* | *SRR6456420* | *SRR6872534* | *SRR6872539* | *SRR8992309* | *SRR8992311* |
| | # reads (M) | 23.79 | 23.20 | 3.49 | 39.44 | 15.42 | 11.77 | 5.96 | 5.81 | 18.01 | 7.11 | 14.96 | 15.96 | 8.26 | 7.38 | 14.10 | 13.19 | 21.09 | 18.21 | 36.78 | 32.83 |
| | T. reads (M) | 21.84 | 20.25 | 3.48 | 39.35 | 15.30 | 11.69 | 5.77 | 5.53 | 10.72 | 6.67 | 11.70 | 11.77 | 7.97 | 7.14 | 13.26 | 11.58 | 19.83 | 17.34 | 36.07 | 32.51 |
| | %Ref. mapped | 70.51 | 76.39 | 48.41 | 48.17 | 51.83 | 56.56 | 69.80 | 64.79 | 82.62 | 86.91 | 23.36 | 29.29 | 51.39 | 44.71 | 67.48 | 56.65 | 56.20 | 58.60 | 93.75 | 91.83 |
| BrumiR | Unipath (k) | 91.8 | 94.3 | 63.3 | 479.5 | 73.1 | 53.9 | 43.7 | 48.8 | 43.5 | 77.3 | 174.9 | 168.0 | 66;9 | 61.3 | 290.1 | 187.2 | 506.7 | 487.2 | 151.2 | 148.4 |
| | candidates | 966 | 1046 | 813 | 2954 | 847 | 743 | 569 | 579 | 824 | 696 | 1501 | 1299 | 1628 | 1350 | 2435 | 1301 | 3992 | 3405 | 899 | 1282 |
| | TP | 401 | 438 | 175 | 497 | 190 | 190 | 230 | 237 | 212 | 192 | 156 | 149 | 228 | 212 | 265 | 164 | 331 | 319 | 123 | 169 |
| miRDeep2 / miR-PREFeR | candidates | 579 | 711 | 278 | 383 | 120 | 115 | 537 | 579 | 171 | 196 | 4401 | 3158 | 4769 | 3798 | 5569 | 3774 | 3257 | 3001 | 311 | 441 |
| | TP | 378 | 428 | 245 | 315 | 86 | 90 | 255 | 264 | 123 | 124 | 376 | 301 | 588 | 507 | 637 | 456 | 975 | 812 | 98 | 115 |
| mirnovo | candidates | 789 | 1017 | 358 | 785 | 5476 | 4181 | 537 | 597 | 955 | 2642 | 91 | 82 | 90 | 78 | 127 | 147 | 230 | 210 | 120 | 134 |
| | TP | 196 | 556 | 234 | 166 | 439 | 365 | 255 | 260 | 100 | 331 | 25 | 11 | 39 | 7 | 37 | 27 | 56 | 67 | 29 | 35 |
| BrumiR | precision | 0.42 | 0.42 | 0.22 | 0.17 | 0.22 | 0.26 | 0.40 | 0.41 | 0.26 | 0.28 | 0.10 | 0.11 | 0.14 | 0.16 | 0.11 | 0.13 | 0.08 | 0.09 | 0.14 | 0.13 |
| | recall | 0.73 | 0.66 | 0.55 | 0.78 | 0.60 | 0.54 | 0.74 | 0.75 | 0.86 | 0.72 | 0.93 | 0.87 | 0.57 | 0.61 | 0.86 | 0.88 | 0.80 | 0.77 | 0.83 | 0.87 |
| | F-score | 0.53 | 0.51 | 0.31 | 0.28 | 0.33 | 0.35 | 0.52 | 0.53 | 0.40 | 0.40 | 0.19 | 0.20 | 0.22 | 0.25 | 0.19 | 0.22 | 0.15 | 0.17 | 0.24 | 0.23 |
| miRDeep2 / miR-PREFeR | precision | 0.65 | 0.60 | 0.88 | 0.82 | 0.72 | 0.78 | 0.47 | 0.46 | 0.72 | 0.63 | 0.09 | 0.10 | 0.12 | 0.13 | 0.11 | 0.12 | 0.30 | 0.27 | 0.32 | 0.26 |
| | recall | 0.38 | 0.36 | 0.32 | 0.31 | 0.32 | 0.30 | 0.31 | 0.36 | 0.14 | 0.10 | 0.78 | 0.86 | 0.68 | 0.75 | 0.70 | 0.58 | 0.77 | 0.80 | 0.83 | 0.79 |
| | F-score | 0.48 | 0.45 | 0.47 | 0.45 | 0.45 | 0.43 | 0.38 | 0.40 | 0.24 | 0.17 | 0.15 | 0.17 | 0.21 | 0.23 | 0.20 | 0.20 | 0.43 | 0.40 | 0.46 | 0.39 |
| mirnovo | precision | 0.25 | 0.55 | 0.65 | 0.21 | 0.08 | 0.09 | 0.47 | 0.44 | 0.10 | 0.13 | 0.27 | 0.13 | 0.43 | 0.09 | 0.29 | 0.18 | 0.24 | 0.32 | 0.24 | 0.26 |
| | recall | 0.23 | 0.71 | 0.69 | 0.22 | 0.75 | 0.77 | 0.77 | 0.76 | 0.38 | 0.69 | 0.43 | 0.17 | 0.25 | 0.00 | 0.38 | 0.34 | 0.50 | 0.66 | 0.55 | 0.55 |
| | F-score | 0.24 | 0.62 | 0.67 | 0.21 | 0.14 | 0.16 | 0.59 | 0.55 | 0.16 | 0.21 | 0.34 | 0.15 | 0.32 | 0.01 | 0.33 | 0.24 | 0.33 | 0.43 | 0.34 | 0.35 |
| BrumiR2ref | precision | 0.44 | 0.44 | 0.21 | 0.18 | 0.28 | 0.30 | 0.38 | 0.40 | 0.41 | 0.46 | 0.11 | 0.11 | 0.14 | 0.15 | 0.11 | 0.12 | 0.09 | 0.09 | 0.17 | 0.16 |
| | recall | 0.60 | 0.51 | 0.42 | 0.71 | 0.45 | 0.38 | 0.58 | 0.56 | 0.63 | 0.41 | 0.90 | 0.84 | 0.47 | 0.54 | 0.76 | 0.76 | 0.72 | 0.72 | 0.81 | 0.84 |
| | F-score | 0.51 | 0.47 | 0.29 | 0.28 | 0.35 | 0.34 | 0.46 | 0.47 | 0.50 | 0.44 | 0.19 | 0.20 | 0.21 | 0.24 | 0.19 | 0.20 | 0.16 | 0.17 | 0.28 | 0.27 |

Table S3: Total elapsed time per tool (seconds) on synthetic datasets. The total elapsed time reported include only the core step of each algorithm.

| | Species | dataset | BrumiR | miRDeep2/miR-PREFeR |
|---|---|---|---|---|
| animal | *Homo sapiens* | f1 | 111 | 6394 |
| | | f2 | 433 | 9661 |
| | *Mus musculus* | f1 | 97 | 5101 |
| | | f2 | 383 | 8169 |
| | *Drosophila melagonaster* | f1 | 25 | 922 |
| | | f2 | 61 | 1182 |
| | *Danio rerio* | f1 | 22 | 1679 |
| | | f2 | 55 | 2067 |
| | *Caenorhabditis elegans* | f1 | 16 | 917 |
| | | f2 | 38 | 1044 |
| plant | *Zea mays* | f1 | 20 | 177 |
| | | f2 | 58 | 173 |
| | *Physcomitrella patens* | f1 | 19 | 196 |
| | | f2 | 55 | 222 |
| | *Oryza sativa* | f1 | 63 | 1661 |
| | | f2 | 227 | 1342 |
| | *Solanum lycopersicum* | f1 | 12 | 1204 |
| | | f2 | 20 | 1193 |
| | *Arabidopsis thaliana* | f1 | 31 | 363 |
| | | f2 | 106 | 362 |

Table S4: Total elapsed time per tool (seconds) on real datasets. The total elapsed time reported include only the core step of each algorithm.

| | Species | SRA_ID | BrumiR | miRDeep2/miR-PREFeR |
|---|---|---|---|---|
| animal | *Homo sapiens* | ERR2403205 | 48 | 6319 |
| | | ERR2403208 | 50 | 5421 |
| | *Mus musculus* | SRR1734814 | 23 | 8342 |
| | | SRR1734817 | 333 | 11103 |
| | *Drosophila melagonaster* | ERR3276595 | 31 | 5382 |
| | | ERR3276596 | 27 | 8270 |
| | *Danio rerio* | SRR2127900 | 20 | 4921 |
| | | SRR2127901 | 22 | 5554 |
| | *Caenorhabditis elegans* | ERR562747 | 24 | 10159 |
| | | ERR562743 | 28 | 8172 |
| plant | *Zea mays* | SRR7801904 | 59 | 52875 |
| | | SRR7801905 | 50 | 13002 |
| | *Physcomitrella patens* | SRR1842130 | 37 | 25909 |
| | | SRR1842134 | 30 | 39644 |
| | *Oryza sativa* | SRR6456419 | 124 | 34346 |
| | | SRR6456420 | 66 | 17916 |
| | *Solanum lycopersicum* | SRR6872534 | 378 | 31902 |
| | | SRR6872539 | 334 | 37629 |
| | *Arabidopsis thaliana* | SRR8992309 | 50 | 4045 |
| | | SRR8992311 | 49 | 1802 |

Table S5: miRNA discovery from the root samples of *Arabidopsis thaliana* using BrumiR.

| | | Raw reads (M) | Processed reads (M) | Unipaths | Candidates | Hairpin validated | Core predictions | Known miRNAs | Putative novel miRNAs |
|---|---|---|---|---|---|---|---|---|---|
| day 5 | 1 | 22.19 | 2.10 | 16,122 | 117 | 72 | 45 | 36 | 5 |
| | 2 | 25.31 | 1.35 | 7,077 | 48 | 34 | | | |
| | 3 | 25.77 | 2.97 | 18,687 | 116 | 69 | | | |
| day 9 | 4 | 24.65 | 15.60 | 90,588 | 989 | 592 | 295 | 78 | 141 |
| | 5 | 21.70 | 14.58 | 61,374 | 685 | 394 | | | |
| | 6 | 24.66 | 14.46 | 107,404 | 1,698 | 1,045 | | | |
| day 13 | 7 | 29.30 | 19.63 | 94,222 | 1,153 | 583 | 468 | 88 | 258 |
| | 8 | 25.73 | 14.49 | 96,656 | 1,238 | 757 | | | |
| | 9 | 27.94 | 20.22 | 144,949 | 1,944 | 1,221 | | | |
| day 17 | 10 | 21.77 | 13.44 | 61,014 | 738 | 441 | 238 | 86 | 96 |
| | 11 | 19.78 | 10.33 | 36,673 | 450 | 245 | | | |
| | 12 | 16.49 | 9.55 | 89,471 | 1,264 | 790 | | | |
| day 21 | 13 | 24.42 | 17.80 | 127,221 | 1,852 | 1,135 | 212 | 89 | 80 |
| | 14 | 16.81 | 6.90 | 34,903 | 480 | 312 | | | |
| | 15 | 22.54 | 5.54 | 26,271 | 434 | 267 | | | |
| day 25 | 16 | 24.19 | 16.31 | 133,004 | 2,069 | 1,273 | 1136 | 133 | 622 |
| | 17 | 35.56 | 27.06 | 181,169 | 2,653 | 1,598 | | | |
| | 18 | 26.05 | 18.21 | 147,944 | 2,245 | 1,379 | | | |

Table S6: Novel miRNAs in the root samples of *Arabidopsis thaliana* predicted by BrumiR.

| **miRID** | **chr:pos** | **mature sequence** | **precursor sequence** |
|---|---|---|---|
| miR-1 | chr5:10807602-10807763 | ACCAAAAACGAAACATTCCCC | TTTATCTGTTAATTTCGTTAGGGGCAATTTTTCGTTTTTGGTGTGGGTATTTTGCATCAATTG GAGTGAGTAGAAGGAGAGGATTGATTGATTTGGTGTTCCAATTCTACCAACCGGAAAGGAT TAGAAGCGATGATGTATCTTCAGACCAACTATTACAT |
| miR-3 | chr3:9240992-9241104 | GGATGAAAGGTTTGACTAGAACT | AATAAATTGGATTTTTTAGTTAGAAAGGTTTGGCAGGACGTTATTTACTAAAAAATAAATGA GTTTTTTAGGATGAAAGGTTTGACTAGAACTGAAGATTTATGTTTATTAT |
| miR-8 | chr1:29612248-29612361 | ATTATGGACCGTCCAACTTGGCCC | TGGGCTGACCATGGACTTGCCCATATGGACATGGTCTTTATTGGGCATGGACATTTTCGGAC CATTGTCCATTATGGACCGTCCAACTTGGCCCATAAAAAAAACTGTCCGTA |

Table S7: Novel microRNAs and their putative interactions obtained using psRNATarget. miRNA_Acc.: microRNA identification; Target_Acc.: mRNA target identification, linked to the *Arabidopsis thaliana* mRNA library with the Araport V11 genome annotation. Expectation: mismatches penalty between mature small RNA and the target sequence, the lower the value the better the prediction (with 5.0 as a maximum threshold). Inhibition: refers to the possible mechanisms used by the sRNA to regulate its mRNA target, described in plants. Target_Desc: refers to the gene description for the mRNA target, found in the Araport V11 annotation. Multiplicity: indicates how many times a sRNA has a target sequence in a unique mRNA.

| **miRNA_Acc.** | **Target_Acc.** | **Expectation** | **Inhibition** | **Target_Desc.** | **Multiplicity** |
|---|---|---|---|---|---|
| miR-1 | AT1G66000.1 | 2.0 | Cleavage | hypothetical protein (DUF577) | 1 |
| miR-1 | AT2G30700.1 | 2.0 | Cleavage | GPI-anchored protein | 1 |
| miR-1 | AT4G16250.1 | 3.0 | Cleavage | phytochrome D | 1 |
| miR-1 | AT4G24740.5 | 3.0 | Cleavage | LAMMER-type protein kinase AFC2 | 1 |
| miR-1 | AT5G03670.2 | 3.5 | Cleavage | histone-lysine N-methyltransferase SETD1B-like protein | 1 |
| miR-1 | AT3G04450.1 | 3.5 | Cleavage | Homeodomain-like superfamily protein | 1 |
| miR-1 | AT4G19920.1 | 3.5 | Cleavage | Toll-Interleukin-Resistance (TIR) domain family protein | 1 |
| miR-1 | AT2G10608.1 | 3.5 | Cleavage | transmembrane protein | 1 |
| miR-1 | AT2G32680.1 | 3.5 | Cleavage | receptor like protein 23 | 1 |
| miR-1 | AT3G18480.1 | 3.5 | Cleavage | CCAAT-displacement protein alternatively spliced product | 1 |
| miR-1 | AT1G14630.2 | 3.5 | Cleavage | XRI1-like protein | 1 |
| miR-1 | AT1G70590.1 | 3.5 | Cleavage | F-box family protein | 1 |
| miR-1 | AT3G21870.1 | 3.5 | Cleavage | cyclin | 1 |
| miR-1 | AT5G49100.1 | 3.5 | Translation | vitellogenin-like protein | 1 |
| miR-3 | AT5G66950.1 | 2.0 | Cleavage | Pyridoxal phosphate (PLP)-dependent transferases superfamily protein | 1 |
| miR-3 | AT2G18720.3 | 2.5 | Cleavage | Translation elongation factor EF1A/initiation factor IF2gamma family protein | 1 |
| miR-3 | AT1G56050.1 | 2.5 | Translation | GTP-binding protein-like protein | 1 |
| miR-3 | AT2G18720.2 | 2.5 | Cleavage | Translation elongation factor EF1A/initiation factor IF2gamma family protein | 1 |
| miR-3 | AT2G18720.1 | 2.5 | Cleavage | Translation elongation factor EF1A/initiation factor IF2gamma family protein | 1 |
| miR-3 | AT3G07540.1 | 2.5 | Cleavage | Actin-binding FH2 (formin homology 2) family protein | 1 |
| miR-3 | AT3G08780.1 | 3.0 | Cleavage | BRISC complex subunit Abro1-like protein | 1 |
| miR-3 | AT4G34920.1 | 3.0 | Cleavage | PLC-like phosphodiesterases superfamily protein | 1 |
| miR-3 | AT2G38060.1 | 3.0 | Cleavage | phosphate transporter | 1 |
| miR-3 | AT3G56080.1 | 3.0 | Cleavage | S-adenosyl-L-methionine-dependent methyltransferases superfamily protein | 1 |
| miR-3 | AT2G41860.1 | 3.0 | Cleavage | calcium-dependent protein kinase 14 | 1 |
| miR-3 | AT4G17570.2 | 3.0 | Cleavage | GATA transcription factor 26 | 1 |
| miR-3 | AT4G35620.1 | 3.5 | Cleavage | Cyclin | 1 |
| miR-3 | AT5G22160.1 | 3.5 | Cleavage | transmembrane protein | 1 |
| miR-3 | AT5G49870.2 | 3.5 | Cleavage | Mannose-binding lectin superfamily protein | 1 |
| miR-3 | AT2G26770.3 | 3.5 | Cleavage | plectin-like protein | 1 |
| miR-8 | AT4G25100.4 | 2.5 | Cleavage | Fe superoxide dismutase 1 | 1 |
| miR-8 | AT5G61630.1 | 3.0 | Cleavage | transmembrane protein | 1 |
| miR-8 | AT5G54020.1 | 3.5 | Cleavage | Cysteine/Histidine-rich C1 domain family protein | 2 |
| miR-8 | AT2G39040.1 | 3.5 | Cleavage | Peroxidase superfamily protein | 1 |
| miR-8 | AT1G04840.1 | 3.5 | Cleavage | Tetratricopeptide repeat (TPR)-like superfamily protein | 1 |
| miR-8 | AT3G20010.8 | 3.5 | Cleavage | SNF2 domain-containing protein / helicase domain-containing protein / zinc finger protein-like protein | 1 |
| miR-8 | AT3G53570.5 | 3.5 | Cleavage | serine/threonine-protein kinase AFC1 | 1 |
| miR-8 | AT1G10580.1 | 3.5 | Cleavage | Transducin/WD40 repeat-like superfamily protein | 1 |
| miR-8 | AT2G02070.2 | 3.5 | Cleavage | indeterminate(ID)-domain 5 | 1 |

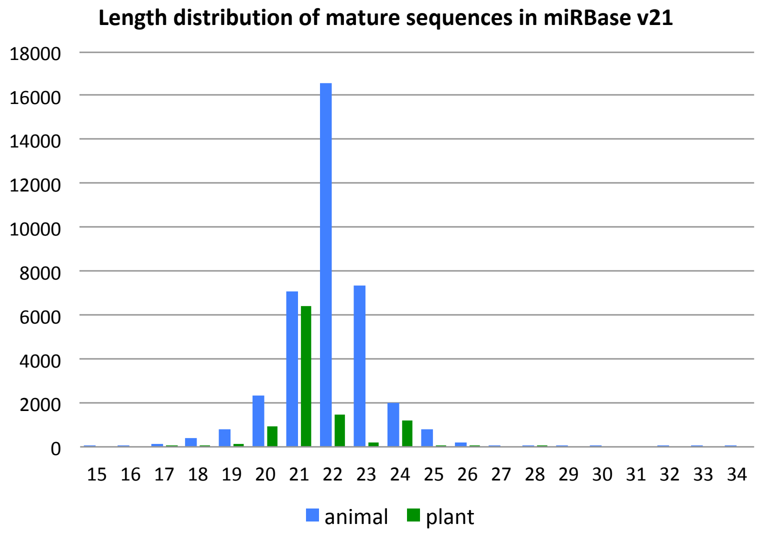

Figure S1: Length distribution of mature sequences in miRBase. We used miRBase v21 with a total of 35828 entries and we observed that the length is between 19-24 nt.

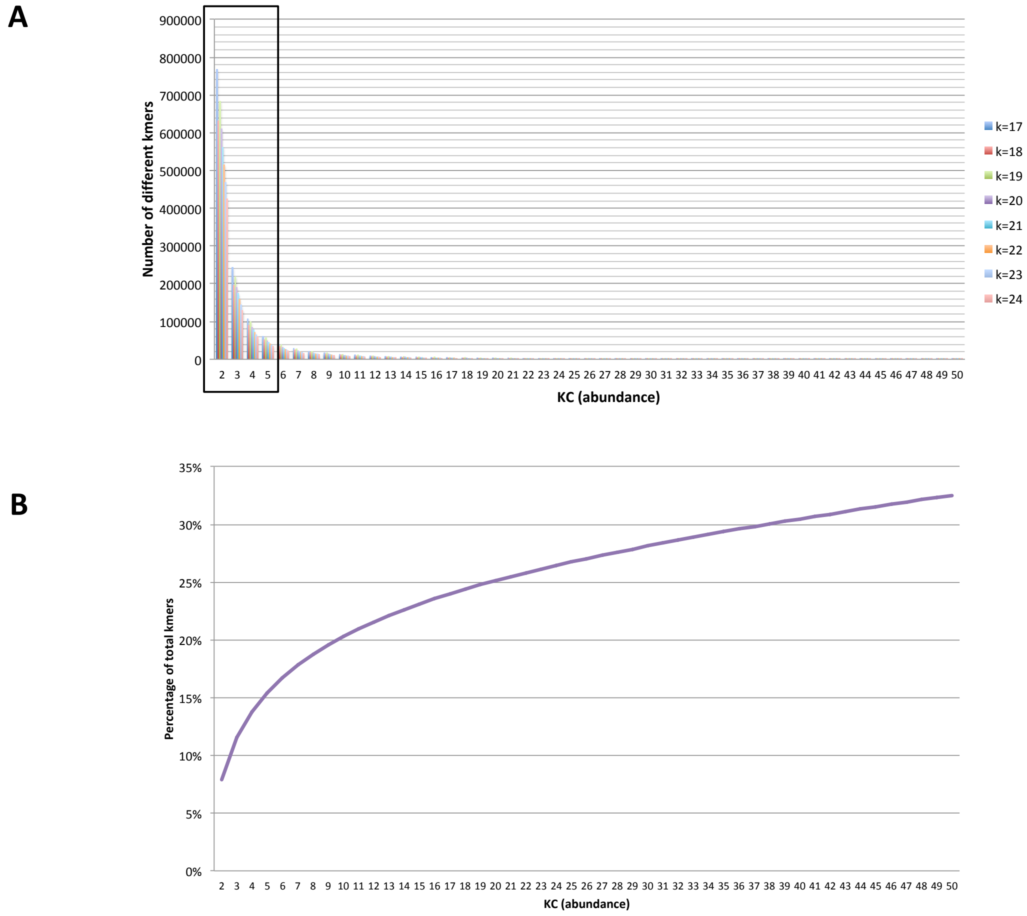

Figure S2: Kmer spectrum of sRNA-seq data. A) The histogram shows the number of distinct kmers (Y-axis) as a function of the read coverage (KC X-axis). In the lower coverage of the spectrum (black rectangle), we observe a high number of distinct kmers which are likely sequencing errors. The kmers that correspond to noise represent approximately less than 15% of the total number of kmers.

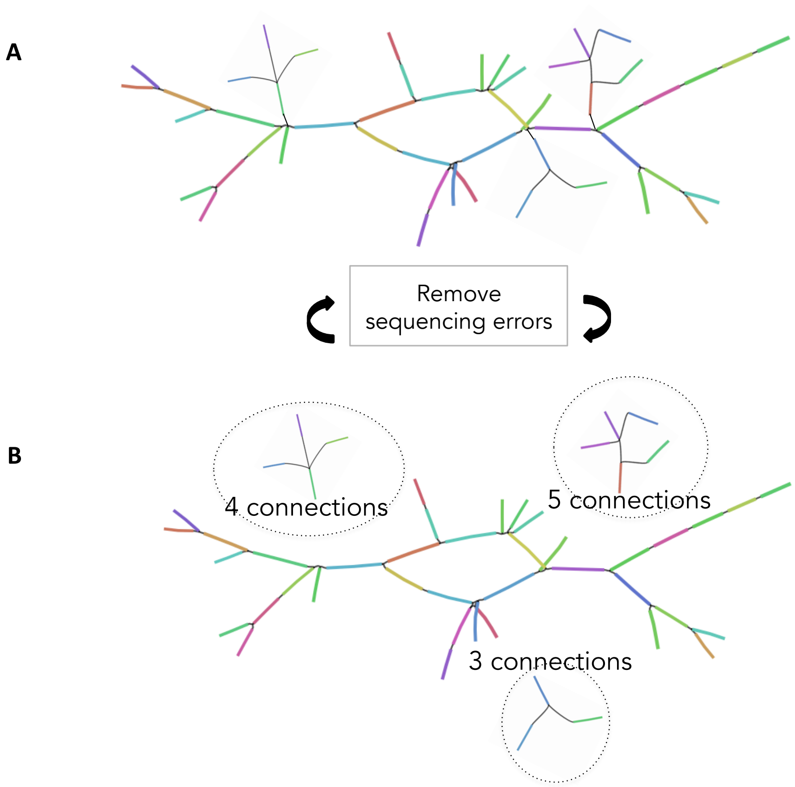

Figure S3: BrumiR classifies low abundance non-linear topologies as sequencing errors. A) BrumiR identifies these topologies connected to the principal structures in the graph, which appear after the first tip removal steps of BrumiR. B) These topologies have low abundance (KM value) and are composed of branching nodes with 3, 4, or 5 connections.

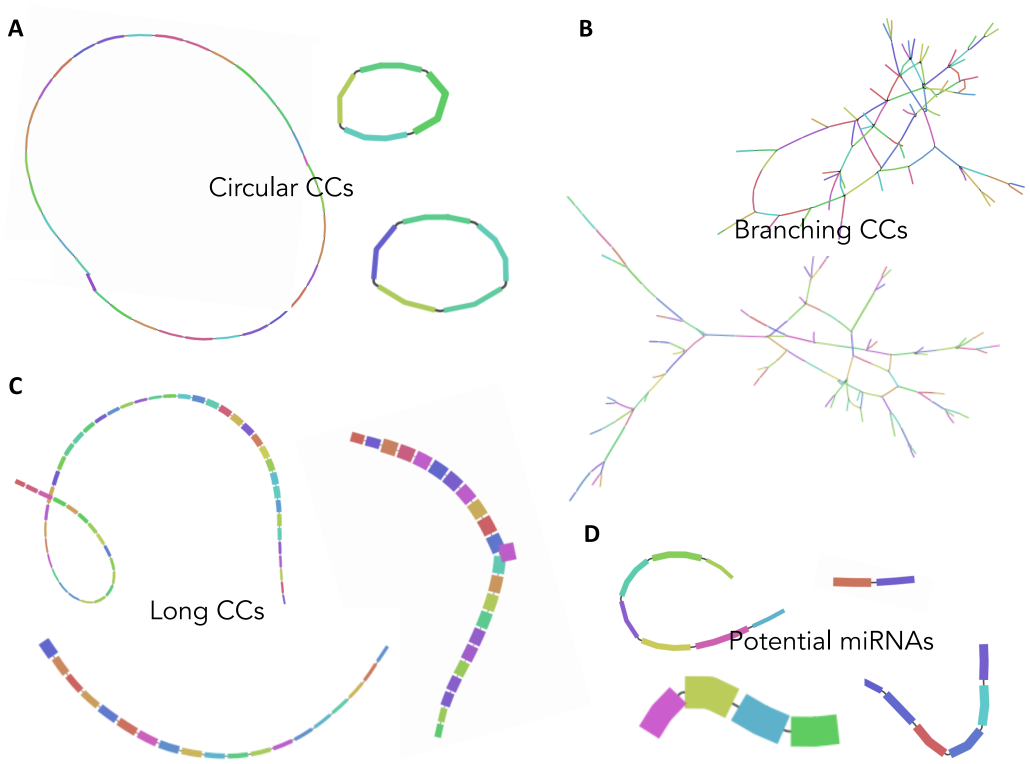

Figure S4: Re-assembling unipaths within each CC. BrumiR re-assembles all unipaths present in a linear CC by bundling the nodes with in and out degree equal to 1 into a new unipath. BrumiR rebuilds each unipath within a CC and classifies them into different types. A) Circular CCs: when all unipaths are have an in and out connection, we classify the CC as a circular sequence that is not a putative miRNA. B) Branching CCs: when we detect a CC with a high number of branching nodes, we do not consider it anymore for the moment, because we consider it related to sequencing errors (usually they have a low KM value). C) Long CCs: when we detect more than 10 unipaths, we can classify them as longer non coding sequences, but we still keep them for later analysis. D) Potential miRNAs: all assembled unipaths (CCs) having a length between 18 and 24 are stored as potential miRNA sequences.

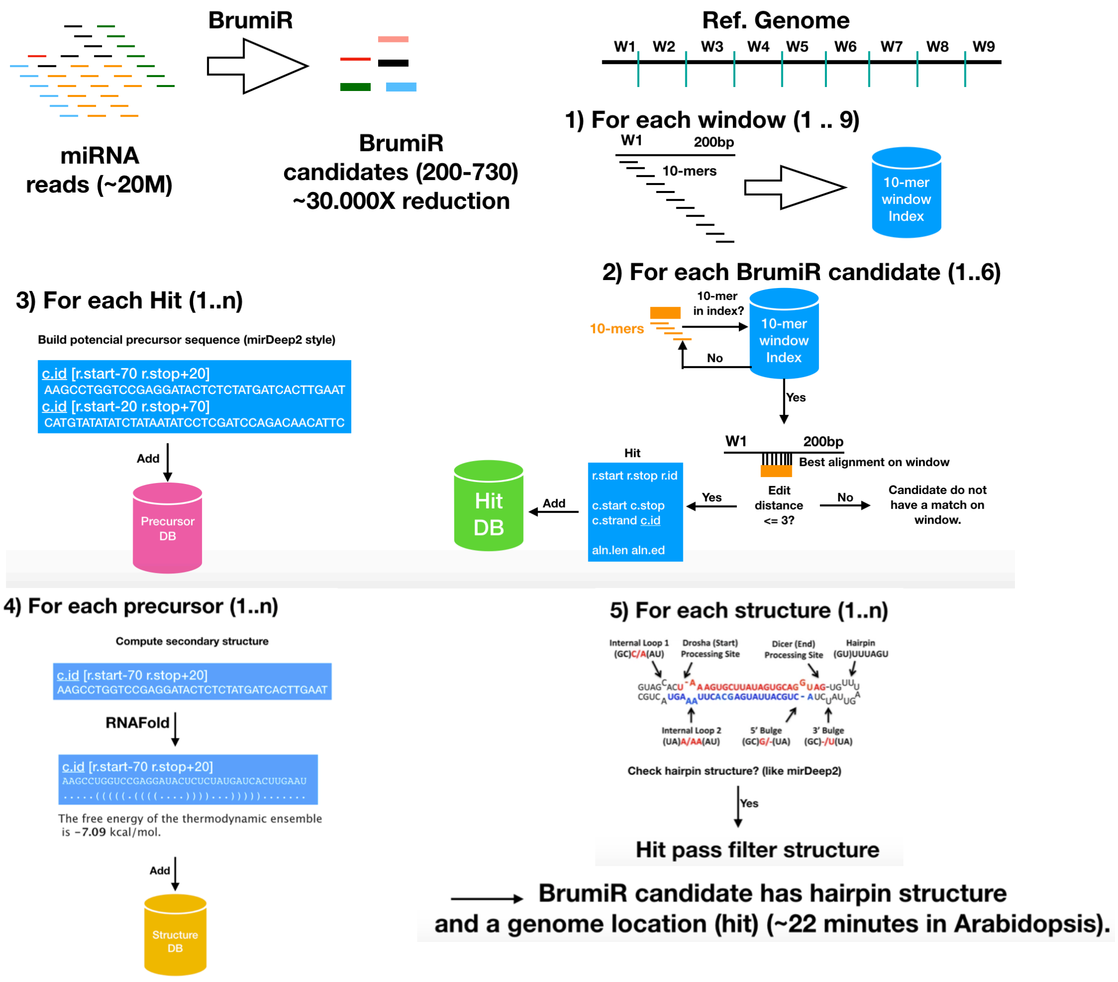

Figure S5: Workflow of the BrumiR2Reference tool. The main steps involved the mapping of the miRNA candidates to the genome using non-overlapping windows (1); each hit is further refined using an exhaustive alignment (2). For each hit, a precursor sequence is built (3), and its secondary structure is determined using RNAfold (4). Finally, structures fulfilling a set of criteria (5) are classified as precursor sequences.

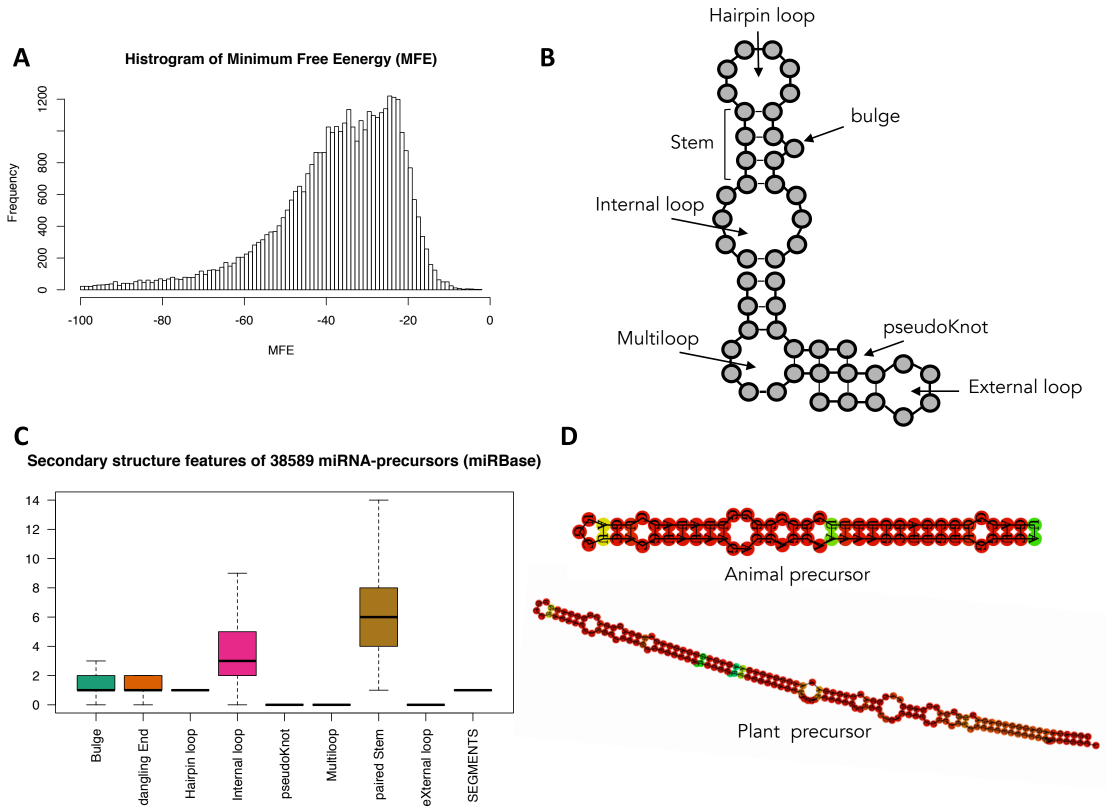

Figure S6: Structure properties of miRBase precursor sequences. A) Free-energy distribution of 38.589 precursor sequences folded with RNAfold. B) Different types of RNA secondary structure elements composing precursor miRNA sequences. C) Analysis of secondary structure elements performed on 38.589 precursor sequences in miRBase using the bpRNA package. D) Examples of precursor sequences for animal and plant species.

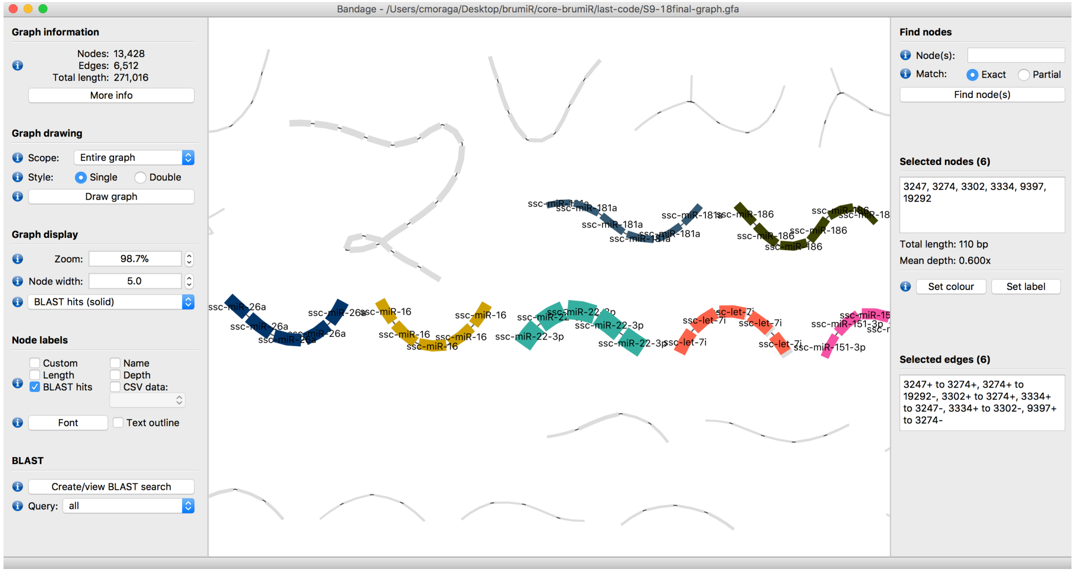

Figure S7: Visualization with Bandage. BrumiR provides an output compatible with the Bandage software, which can be employed to visualize and explore the results in a user-friendly way.

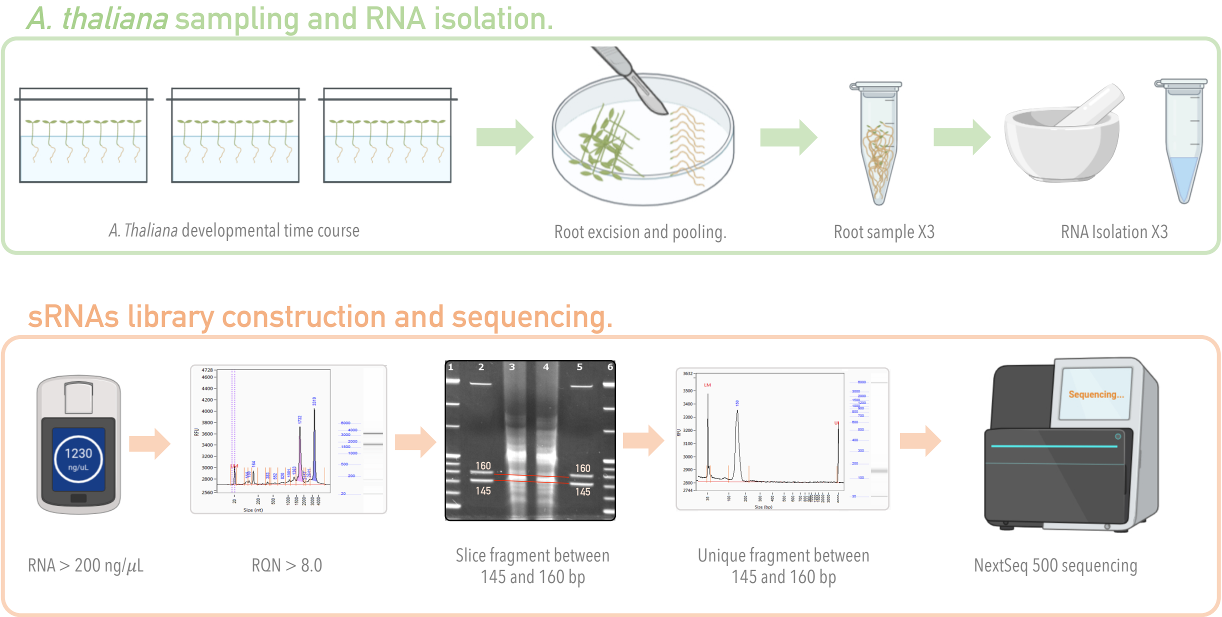

Figure S8: Experimental procedure of the *Arabidopsis thaliana* roots sampling and the sRNA-seq libraries construction. The seedlings for each sampling point are excised from their aerial shoots, the roots are pooled and stored in triplicates for RNA isolation. The RNAs are evaluated for concentration and integrity, above 200ng/uL, and an RQN score over 8.0 respectively, to begin a sRNA-Seq library construction. The 145-160 bp library was purified from polyacrylamide gels and validated as a unique fragment between 145-160 bp. Finally, the successful libraries were processed for next-generation sequencing (NGS) procedures. Representative AATI Fragment Analyzer electropherograms are shown for RNA integrity and sequencing library validation; LM, a lower marker at 20 nt and 35 bp; UM: upper marker at 4,000 nt and 6,000 bp. Polyacrylamide gel electrophoresis (PAGE) of reverse-transcribed cDNAs from small RNAs are shown. Red lines indicate fragments of interest (including sRNAs and miRNAs). Extraction procedures for fragments from PAGE are described in the library construction procedures by the manufacturer. * Figure partially created with BioRender.com.

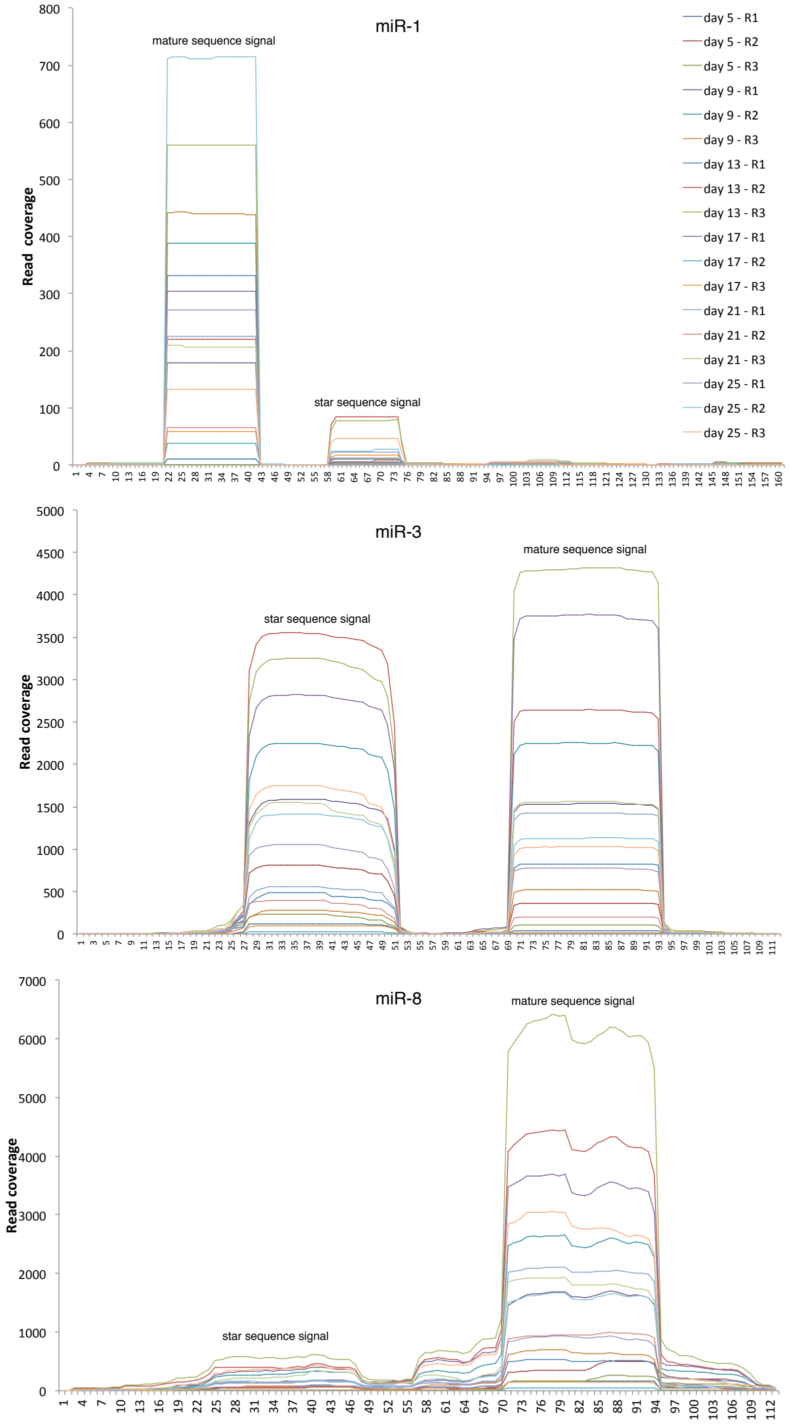

Figure S9: Read coverage of novel precursor candidates found in the roots of *Arabidopsis thaliana*. Reads of each experimental condition were mapped back to the novel precursor sequences using Bowtie. Samtools was used to compute the read coverage (depth) along each precursor sequence. The three novel precursor sequences show the read signature peaks.
